# Alzheimer-related pathogenesis is dependent on neuronal receptor PTPσ

**DOI:** 10.1101/079806

**Authors:** Yuanzheng Gu, Yaoling Shu, Angela W. Corona, Kui Xu, Allen F. Yi, Shannon Chen, Man Luo, Michel L. Tremblay, Randy J. Nelson, Gary E. Landreth, Jerry Silver, Yingjie Shen

## Abstract

Due to limited understanding of disease mechanisms and the lack of molecular targets, translational research for Alzheimer disease has not been fruitful hitherto. Here we report findings that indicate neuronal receptor phosphatase PTPσ as a potential therapeutic target for this dementia. In two TgAPP mouse models, a spectrum of Alzheimer-related pathologies, including aged-induced progression of β-amyloidosis, Tau aggregation, neuroinflammation, synaptic loss, as well as behavioral deficits, all show unambiguous dependency on PTPσ. APP amyloidogenic metabolites diminish upon PTPσ genetic depletion or pharmacological inhibition. Binding to APP in the brain, PTPσ regulates APP proteolytic metabolism via its phosphatase activity, likely through downstream signaling that modulates APP membrane localization and affinity to the β-secretase, in a specific manner that does not broadly affect β- and γ-secretase processing of other major substrates. Together, these findings unveil a gatekeeping role of PTPσ upstream in Alzheimer-like pathogenic pathway.

---

A definitive pathological hallmark of Alzheimer’s disease (AD) is the progressive aggregation of β-amyloid (Aβ) peptides in the brain, a process also known as β-amyloidosis, which is often accompanied by neuroinflammation and formation of neurofibrillary tangles containing Tau, a microtubule binding protein.

Although the etiological mechanisms of AD have been an ongoing debate, concrete evidence from human genetic studies showed that overproduction of Aβ due to gene mutations inevitably inflicts cascades of cytotoxic events, ultimately leading to neurodegeneration and decay of brain functions. Contrarily, a gene variation that reduces Aβ generation protects against this dementia (*1*). Accumulation of Aβ peptides, especially in their soluble forms, is therefore recognized as a key culprit in the development of AD (*2*). In the brain, Aβ peptides mainly derive from sequential cleavage of neuronal Amyloid Precursor Protein (APP) by the β- and γ-secretases. However, despite decades of research, molecular regulation of the amyloidogenic secretase activities remains poorly understood, hindering the design of therapeutics to specifically target the APP amyloidogenic pathway.

Pharmacological inhibition of the β- and γ-secretase activities, although effective in suppressing Aβ production, interferes with physiological function of the secretases on their other substrates. Such intervention strategies therefore are often innately associated with untoward side effects, including disrupted cognitive functions, which may provide explanation for several failed clinical trials in the past (*3–5*). To date, no therapeutic regimen is available to prevent the onset of AD or curtail its progression.

Here we report our findings that identify neuronal receptor PTPσ (Protein Tyrosine Phosphatase Sigma) as a potential molecular target to curtail APP amyloidogenic metabolism, to a similar extent as the Iceland mutation that protects against this dementia (*1*). Via its phosphatase signaling, PTPσ functions in a specific manner that does not broadly inhibit β- and γ-secretase activities, but likely through modulating APP distribution to the lipid rafts and its affinity to the β-secretase.

Besides Aβ, Tau is another biomarker that has been intensively studied in AD. Cognitive decline in patients often correlates better with Tau pathology than with Aβ burden (*6, 7*). Overwhelming evidence also substantiated that malfunction of Tau contributes to synaptic loss and neuronal deterioration (*8*). However, what triggers the pathological changes of Tau in AD remains a mystery. Whether neurotoxic Aβ can lead to Tau pathology *in vivo* has been a debate since quintessential neurofibrillary Tau tangles have not been reported in any of the APP transgenic mouse models, even in those with severe cerebral β-amyloidosis. We now show that in two APP transgenic mouse models, Tau protein forms small aggregates during aging, with a similar distribution pattern as seen in postmortem AD brains, suggesting that Tau misfolding, although not in typical tangle morphologies, can develop as an event downstream from the overexpression of amyloidogenic APP transgenes. Genetic depletion of PTPσ, which diminishes Aβ levels, also inhibits the aggregation of Tau.

Consistently in the two mouse models we studied, a spectrum of AD-related neuropathologies and behavioral deficits all demonstrate a clear dependency on PTPσ, further supporting a role of this neuronal receptor as a crucial gatekeeper in AD-like pathogenesis.

## PTPσ is an APP binding partner in the brain

Previously identified as a neuronal receptor of extracellular proteoglycans (*9–11*), PTPσ is expressed in adult mammals with a pattern highly enriched in the nervous system, most predominantly in the hippocampus (*12, 13*), one of earliest affected brain regions in AD. Through immunohistochemistry and confocal imaging, we found that PTPσ and APP (the precursor of Aβ) colocalize in hippocampal pyramidal neurons of adult rat brains, most intensively in the initial segments of apical dendrites, and in the perinuclear and axonal regions with a punctate pattern (Fig. 1A-F). To assess whether this colocalization reflects a binding interaction between these two molecules, we tested their co-immunoprecipitation from brain homogenates. In brains of rats and mice with different genetic background, using various antibodies of APP and PTPσ, we consistently detected a fraction of PTPσ that co-immunoprecipitates with APP, providing evidence of a molecular complex between these two transmembrane proteins (Fig. 1H, I; fig. S1).

**Figure 1.**
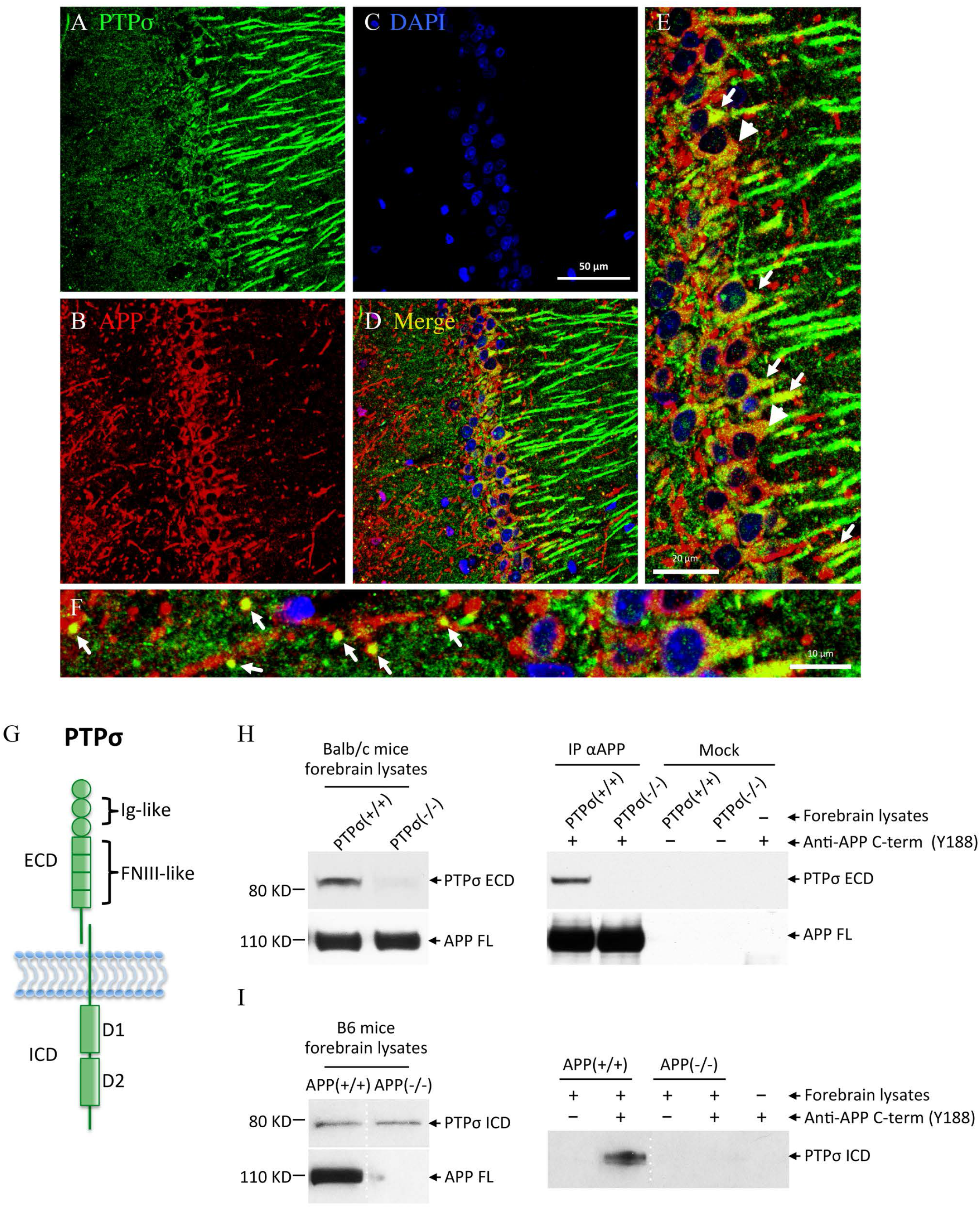
PTPσ is an APP binding partner in the brain. (**A**-**F**) Colocalization of PTPσ (**A**, green) and APP (**B**, red) in hippocampal CA1 neurons of adult rat is shown by confocal imaging. Nuclei are stained with DAPI (**C**, blue). (**D**) Merge of three channels. Scale bar, 50 µm. (**E**) Zoom-in image of the soma layer in **D**. Arrows, intensive colocalization of PTPσ and APP in the initial segments of apical dendrites; arrow heads, punctates of colocalization in the perinuclear regions. Scale bar, 20 µm. (**F**) Zoom-in image of the very fine grained punctates in the axonal compartment in **D**. Arrows points to the colocalization of PTPσ and APP in axons projecting perpendicular to the focal plane. Scale bar, 10 µm. (**G**) Schematic diagram of PTPσ expressed on cell surface as a two-subunit complex. PTPσ is post-translationally processed into an extracellular domain (ECD) and a transmembrane-intracellular domain (ICD). These two subunits associate with each other through noncovalent bond. Ig-like, immunoglobulin-like domains; FNIII-like, fibronectin III-like domains; D1 and D2, two phosphatase domains. (**H**, **I**) Co-immunoprecipitation (co-IP) of PTPσ and APP from mouse forebrain lysates. Left panels, expression of PTPσ and APP in mouse forebrains. Right panels, IP using an antibody (Y188) specific for the C-terminus (C-term) of APP. Full length APP (APP FL) is detected by the same anti-APP C-term antibody. (**H**) PTPσ co-IP with APP from forebrain lysates of wild type but not PTPσ-deficient mice (Balb/c background), detected by an antibody against PTPσ-ECD. (**I**) PTPσ co-IP with APP from forebrain lysates of wild type but not APP knockout mice (B6 background), detected by an antibody against PTPσ-ICD. Dotted lines in **I** indicate lanes on the same western blot exposure that were not adjacent to each other. Images shown are representatives of more than five independent experiments using mice between ages of 1month to 2 years.

## PTPσ depletion diminishes amyloidogenic products of APP

The molecular interaction between PTPσ and APP prompted us to investigate whether PTPσ plays a role in amyloidogenic processing of APP. In neurons, APP is mainly processed through alternative cleavage by either α- or β-secretase. These secretases release the N-terminal portion of APP from its membrane-tethering C-terminal fragment (CTFα or CTFβ, respectively), which can be further processed by the γ-secretase (*14, 15*). Sequential cleavage of APP by the β- and γ-secretases is regarded as amyloidogenic processing since it generates Aβ peptides (*16*). When overproduced, the Aβ peptides can form soluble oligomers that trigger ramification of cytotoxic cascades, whereas progressive aggregation of Aβ eventually results in the formation of senile plaques in the brains of AD patients (Fig. 2A). To test the effect of PTPσ on APP amyloidogenic processing, we analyzed the levels of APP β- and γ-cleavage products in mouse brains with or without PTPσ.

**Figure 2.**
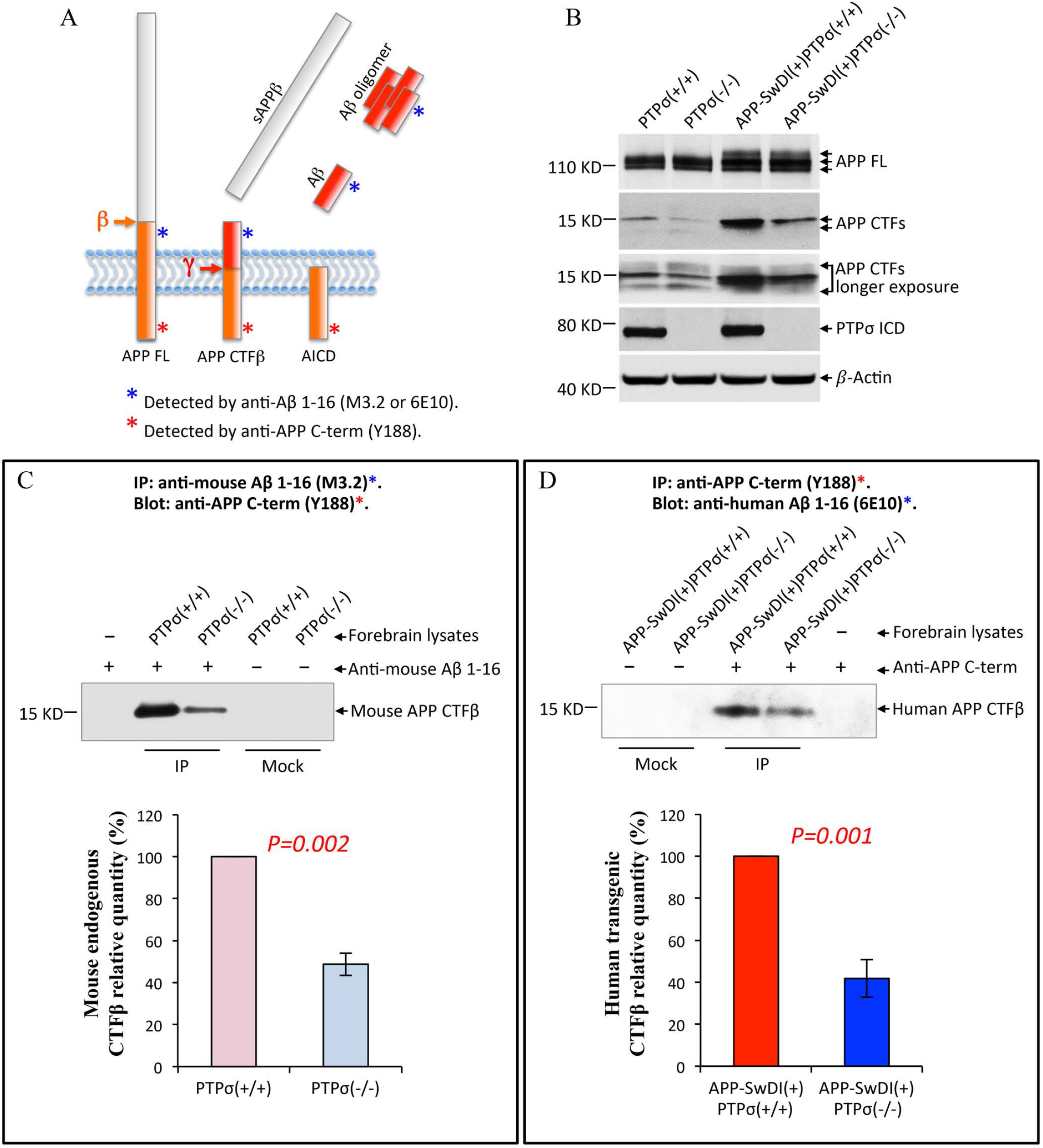
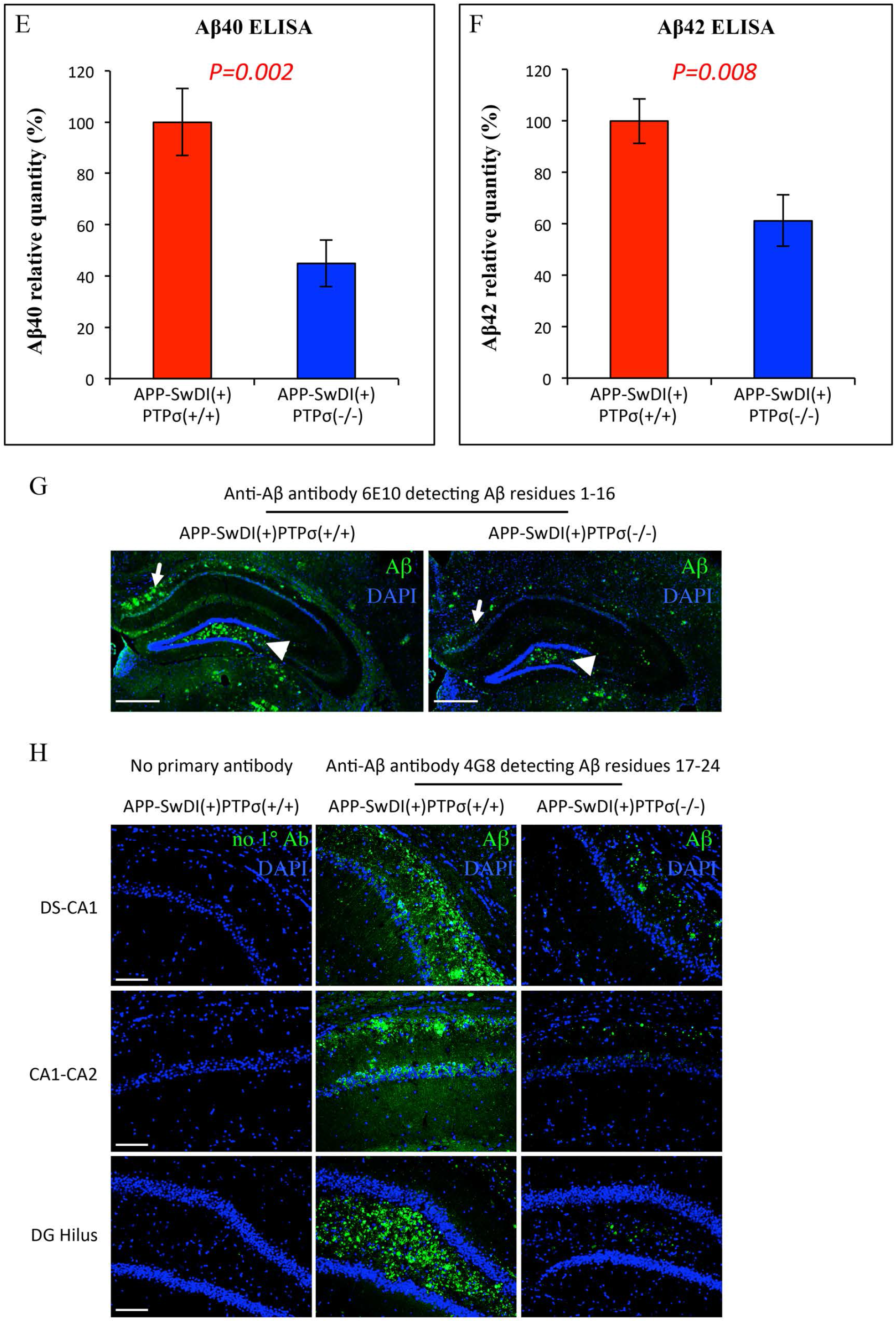
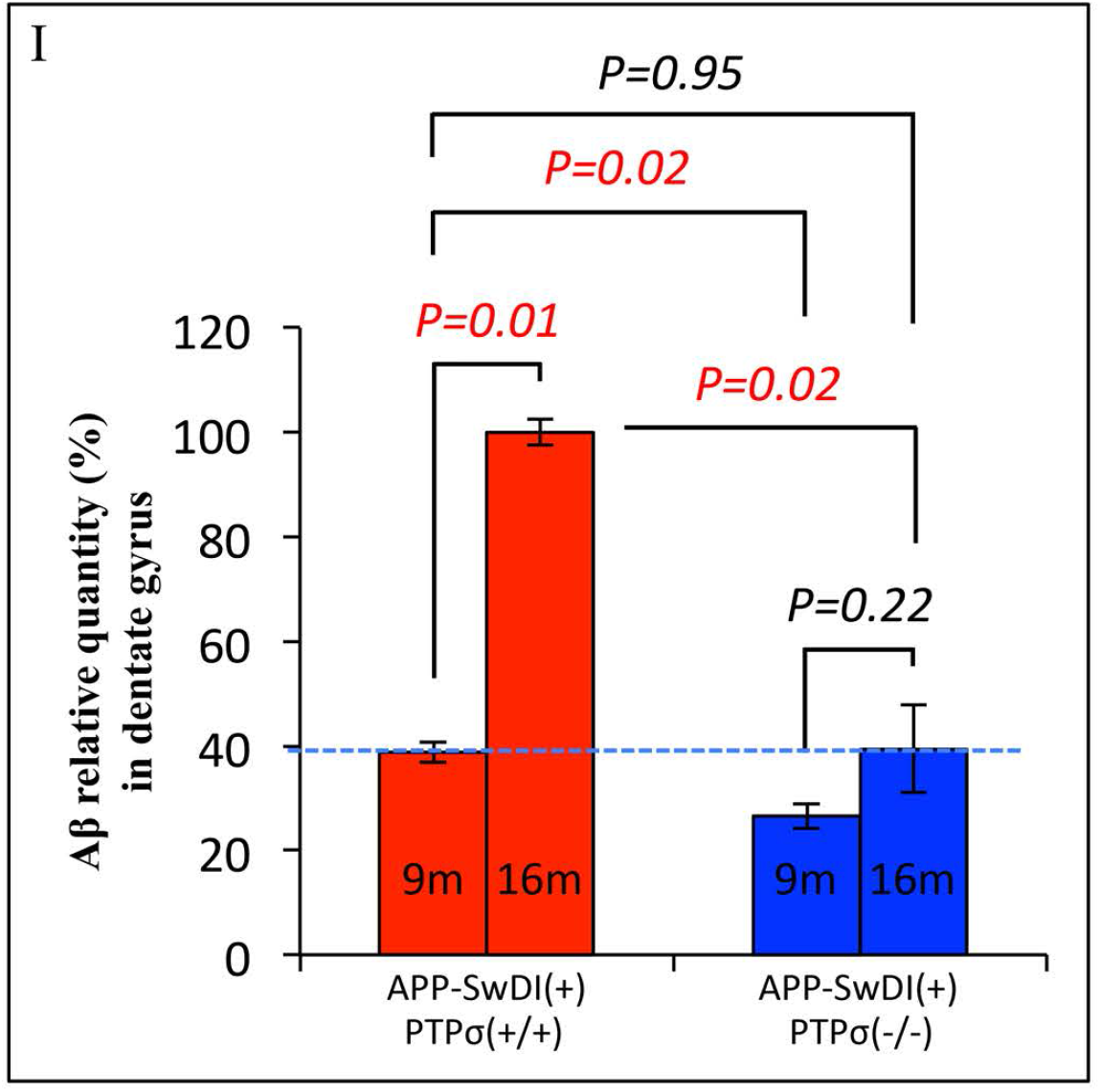
Genetic depletion of PTPσ reduces β-amyloidogenic products of APP. (**A**) Schematic diagram showing amyloidogenic processing of APP by the β- and γ-secretases. Full length APP (APP FL) is cleaved by β-secretase into soluble N-terminal (sAPPβ) and C-terminal (CTFβ) fragments. APP CTFβ can be further processed by γ-secretase into a C-terminal intracellular domain (AICD) and an Aβ peptide. Aggregation of Aβ is a definitive pathology hallmark of AD. (**B**) PTPσ deficiency reduces the level of an APP CTF at about 15 KD in mouse forebrain lysates, without affecting the expression of APP FL. Antibody (Y188) against the C-terminus of APP recognizes APP FL and CTFs of both mouse and human origins. (**C**, **D**) The 15 KD APP CTF is identified as CTFβ by immunoprecipitation (IP) followed with western blot analysis, using a pair of antibodies as marked in the diagram (**A**). Antibodies against amino acids 1-16 of Aβ (anti-Aβ 1-16) detect CTFβ but not CTFα, as the epitope is absent in CTFα. (**C**) Mouse endogenous CTFβ level is reduced in PTPσ-deficient mouse brains. 4 repeated experiments were quantified by densitometry. (**D**) Human transgenic CTFβ level is reduced in PTPσ-deficient mouse brains harboring human APP-SwDI transgene. 6 repeated experiments were quantified by densitometry. Within each experiment in both **C** and **D**, the value from PTPσ deficient sample was normalized to that from the paired wild type sample. (**E, F**) PTPσ deficiency reduces the levels of Aβ40 (**E**) and Aβ42 (**F**) in TgAPP-SwDI mice as measured by ELISA assays. n=12 for each group. The mean values from PTPσ deficient samples was normalized to that from wild type samples. (**G**, **H**) Aβ deposition in the hippocampus of 10-month old TgAPP-SwDI mice. PTPσ deficiency significantly decreases Aβ burden. Aβ (green) is detected by immunofluorescent staining using anti-Aβ antibodies clone 6E10 (**G**) and clone 4G8 (**H**). DAPI staining is shown in blue. (**H**) Upper panels, the stratum oriens layer between dorsal subiculum (DS) and CA1 (also shown with arrows in **G**); middle panels, oriens layer between CA1 and CA2; lower panels, the hilus of dentate gyrus (DG, also shown with arrow heads in **G**). Left column, control staining without primary antibody (1° Ab). No Aβ signal is detected in non-transgenic mice (data not shown). Images shown are representatives from 5 pairs of gender-matched littermates of similar age. Scale bars, 500 µm in **G** and 100 µm in **H**. (**I**) Genetic depletion of PTPσ suppresses the progression of Aβ pathology in TgAPP-SwDI mice. ImageJ quantification of Aβ immunofluorescent staining (with 6E10) in DG hilus from 9- and 16-month old TgAPP-SwDI mice. n=3 for each group. Total integrated density of Aβ in DG hilus was normalized to the area size of the hilus to yield the average intensity as shown in the bar graph. Mean value of each group was normalized to that of 16 month old TgAPP-SwDI mice expressing wild type PTPσ. All *p* values, Student’s *t* test, 2-tailed. All data are mean ± SEM.

Western blot analysis with protein extracts from mouse brains showed that genetic depletion of PTPσ does not affect the expression level of full length APP (Fig. 2B; fig S2A). However, a C-terminal APP fragment, at a molecular weight consistent with that of CTFβ, is found reduced in PTPσ-deficient mice when compared to their wild type littermates (Fig. 2B). Additionally, in the TgAPP-SwDI and TgAPP-SwInd mice, each expressing a human APP transgene harboring the Swedish mutation near the β-cleavage site (*17, 18*), we observed a similar decrease of an APP CTF upon PTPσ depletion (Fig. 2B; fig. S2B). Because the Swedish mutation is prone to β-cleavage, the predominant form of APP CTF in these transgenic mice is predicted to be CTFβ. Hence the reduction of APP CTF in PTPσ-deficient APP transgenic mice may suggest a regulatory role of PTPσ on CTFβ level. However, since the antibody used in these experiments (against APP C-terminus) can recognize both CTFα and CTFβ, as well as their phosphorylated species (longer exposure of western blots showed multiple CTF bands), judging the identity of the reduced CTF simply by its molecular weight may be inadequate. We therefore performed CTFβ immunopurification with subsequent western blot detection, using an antibody that recognizes CTFβ but not CTFα (Fig. 2C, D; fig. S2C, D). The identity of CTFβ is further verified by its negative reactivity to an antibody that detects N-terminal to the β-cleavage site (clone 1G6, data not shown). With this definitive method, we confirmed that PTPσ depletion decreases the level of CTFβ originated from both mouse endogenous and human transgenic APP.

Because CTFβ is an intermediate proteolytic product between β- and γ-cleavage, its decreased steady state level could result from either reduced production by β-cleavage or increased degradation by subsequent γ-secretase cleavage (Fig. 2A). To distinguish between these two possibilities, we additionally measured the level of Aβ peptides, which are downstream products from CTFβ degradation by γ-cleavage. Using ELISA assays with brain homogenates from the TgAPP-SwDI mice, we found that PTPσ depletion decreases the levels of Aβ peptides to a similar degree as that of CTFβ (Fig. 2E, F), reminiscent of the reduction effects by the Iceland (protective) mutation of APP (*1*). Likewise, immunohistological staining showed that PTPσ deficiency also substantially reduces cerebral Aβ deposition during the course of aging, when comparing gender-matched APP transgenic littermates with or without PTPσ (Fig. 2G, H; fig. S2E, F). Thus, the concurrent decrease of β- and γ-cleavage products argues against an increased γ-secretase activity. Although it cannot be ruled out that some alternative uncharacterized pathway may contribute to such parallel decrease, these data strongly support a reduced β-secretase cleavage of APP in PTPσ-deficient brains, which results in lower levels of both CTFβ and downstream Aβ production.

## Progression of β-amyloidosis is curtailed in the absence of PTPσ

Progressive cerebral Aβ deposition (β-amyloidosis) is regarded as a benchmark of AD progression, and this pathological development is observed in many TgAPP mouse models during the course of aging. Given the subsided Aβ levels in the absence of PTPσ, it is conceivable that β-amyloidosis progression would also be deferred. We hence monitored Aβ deposition in the brains of TgAPP-SwDI mice from 9-month to 16-month of ages. At mid-age of 9 to 11 months, Aβ deposits are found predominantly in the hippocampus, especially in the hilus of the dentate gyrus (DG) (Fig. 2G, H). By 16 months (aged), the pathology spreads massively throughout the entire brain. The propagation of Aβ deposition, however, is curbed by genetic depletion of PTPσ, as quantified using the DG hilus as a representative area (Fig. 2I). Between the ages of 9 and 16 months, the Aβ burden is more than doubled in TgAPP-SwDI mice expressing wild type PTPσ [APP-SwDI(+)PTPσ(+/+)] (red bars), but only shows marginal increase in the transgenic mice lacking functional PTPσ [APP-SwDI(+)PTPσ(−/−)] (blue bars). The slow increment of Aβ deposition in the absence of PTPσ and the fact that Aβ loads are similar between 16-month old mice without PTPσ and 9-month old mice with PTPσ (*p*=0.95, dotted line), confirms the attenuation of disease progression by PTPσ deficiency (Fig. 2I).

## PTPσ provides a specific regulation on APP metabolism

The constraining effect of PTPσ on APP amyloidogenic products led us to further question whether this observation reflects a specific regulation of APP metabolism, or alternatively, a general modulation on the β- and γ-secretases. We therefore assessed the expression level of these secretases in mouse brains with or without PTPσ, and found no change for BACE1 (the β-secretase in the brain) or the essential subunits of γ-secretase (Fig. 3A, B). Additionally, we tested whether PTPσ broadly modulates β- and γ-secretase activities, by examining the proteolytic processing of their other substrates. Besides APP, Neuregulin1 (NRG1) (*19–21*) and Notch (*22–24*) are the major *in vivo* substrates of BACE1 and γ-secretase, respectively. Neither BACE1 cleavage of NRG1 nor γ-secretase cleavage of Notch is affected by PTPσ deficiency (Fig. 3C, D). Taken together, these data rule out a generic modulation of β- and γ-secretases, but rather suggest a regulatory specificity on APP metabolism by PTPσ.

**Figure 3.**
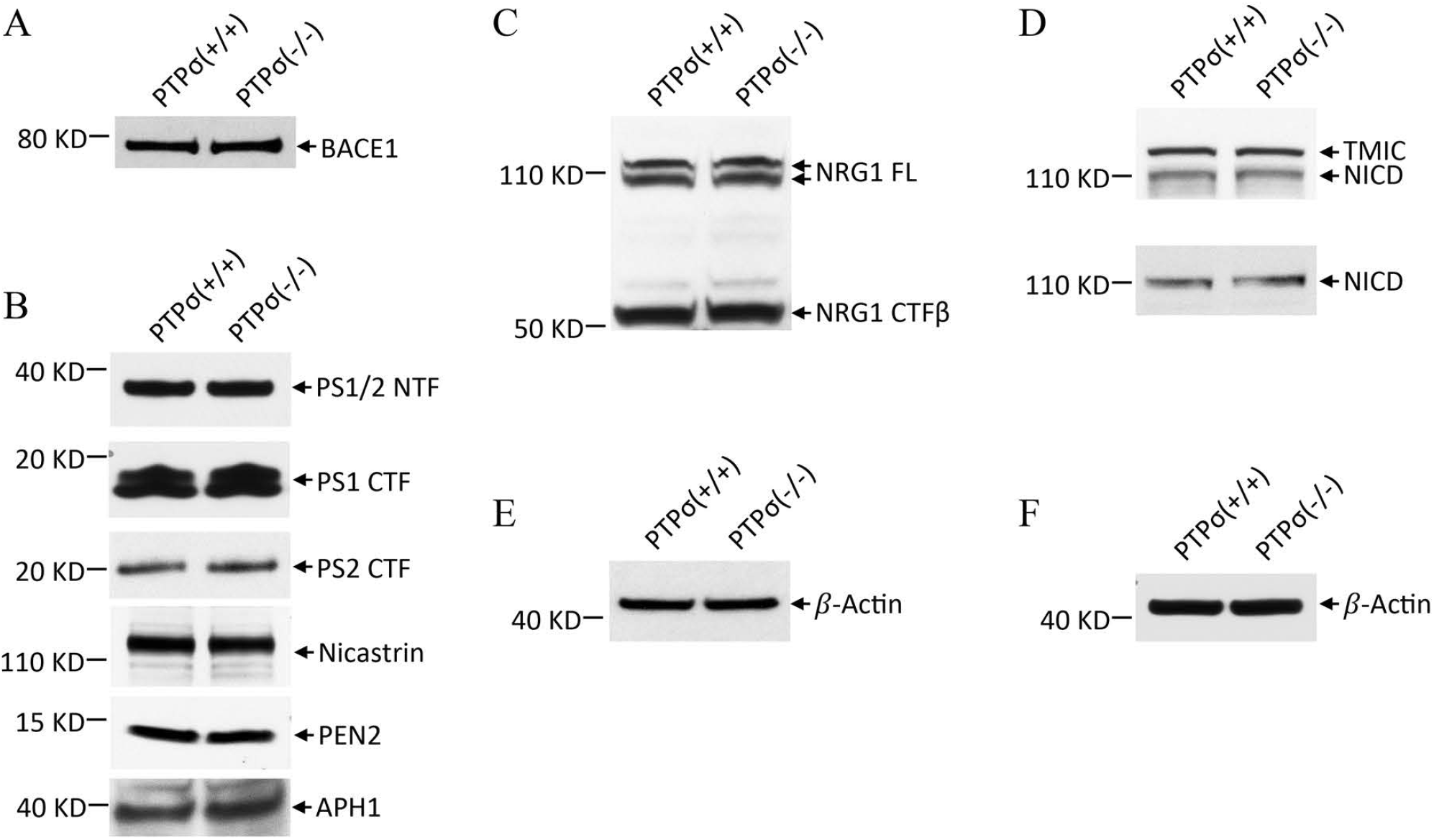
PTPσ does not generically modulate β- and γ-secretases. Neither expression levels of the secretases or their activities on other major substrates are affected by PTPσ depletion. Mouse forebrain lysates with or without PTPσ were analyzed by western blot. (**A, B**) PTPσ deficiency does not change expression level of BACE1 (**A**) or γ-secretase subunits (**B**). Presenilin1 and 2 (PS1/2) are the catalytic subunits of γ-secretase, which are processed into N-terminal and C-terminal fragments (NTF and CTF) in their mature forms. Nicastrin, Presenilin Enhancer 2 (PEN2), and APH1 are other essential subunits of γ-secretase. (**C**) PTPσ deficiency does not change the level of Neuregulin1 (NGR1) CTFβ, the C-terminal cleavage product by BACE1. NRG1 FL, full length Neuregulin1. (**D**) The level of Notch cleavage product by γ-secretase is not affected by PTPσ deficiency. TMIC, Notch transmembrane/intracellular fragment, which can be cleaved by γ-secretase into a C-terminal intracellular domain NICD (detected by an antibody against Notch C-terminus in the upper panel, and by an antibody specific for γ-secretase cleaved NICD in the lower panel). (**E**) β-Actin loading control for **A** and **C**. (**F**) β-Actin loading control for **B** and **D**. All images shown are representatives of at least three independent experiments, each using a pair of age- and gender-matched mice with or without PTPσ deficiency.

It is worth noting that an enzymatic inhibition of the β- or γ-secretase will inevitably suppress the normal functions of these enzymes on a broad array of their substrates. Yet the signaling of NRG1 and Notch, for example, is essential for cognitive housekeeping. Hence approaches of this nature, even though effective in lowering Aβ burden, are innately associated with untoward effects, including disruption of cognitive functions, which may explain the failure to achieve functional efficacy in several Alzheimer clinical trials (*3–5*). The specific regulation on APP metabolism therefore is an important safety feature for PTPσ as a potential therapeutic target.

## PTPσ regulates BACE1-APP affinity in the brain

In an effort to explore the underlying mechanism of PTPσ regulatory effects, particularly its specificity on APP metabolism, we examined the strength of BACE1-APP interaction. The *in vivo* affinity between BACE1 and APP was tested by enzyme-substrate co-immunoprecipitation from mouse brain homogenates in buffers with serially increased detergent stringency. Whereas BACE1-APP association is nearly equal in wild type and PTPσ-deficient brains under mild nonionic buffer conditions, increasing detergent stringency in the buffer unveils that the molecular complex is more vulnerable to dissociation in brains lacking PTPσ (Fig. 4A-C). Such decreased BACE1-APP affinity in PTPσ-deficient brains may indicate a less accessibility of BACE1 to APP and therefore provide a mechanistic basis for the diminished levels of CTFβ and its derivative Aβ.

**Figure 4.**
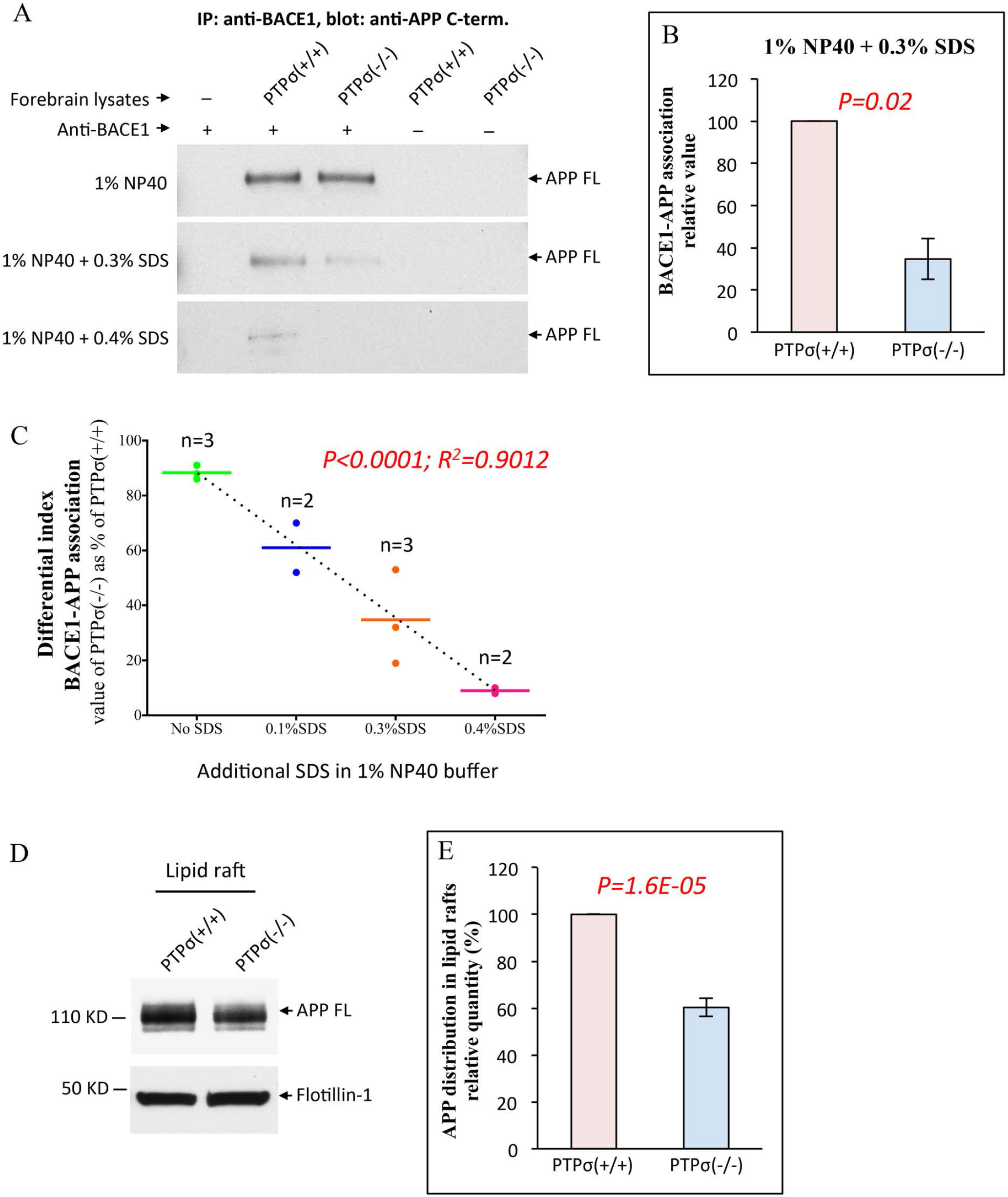
PTPσ deficiency decreases BACE1-APP affinity in the brain and APP distribution in lipid rafts. (**A**) Co-immunoprecipitation experiments show nearly equal BACE1-APP association in wild type and PTPσ-deficient mouse forebrains under mild detergent condition (1% NP40). However, the association in PTPσ-deficient brains is more vulnerable to increased detergent stringency as compared to that in wild type brains. Panels of blots show full length APP (APP FL) pulled down with an anti-BACE1 antibody from mouse forebrain lysates. NP40, Nonidet P-40, non-ionic detergent. SDS, Sodium dodecyl sulfate, ionic detergent. (**B**) Co-immunoprecipitation under buffer condition with 1% NP40 and 0.3% SDS, as shown in the middle panel of **A**, were repeated with three pair of mice. Results were quantified by densitometry, and within each experiment the value from PTPσ-deficient sample was calculated as a percentage of the paired wild type sample value (also shown in orange in **C**). (**C**) Regression analysis of data points from 10 pairs age- and gender-matched mice, each with 1 wild type and 1 PTPσ-deficient. The percentage value (differential index) from each experiment was derived using the same method described for **B** and plotted as a single data point. Increasingly stringent buffer conditions manifest a lower BACE1-APP affinity in PTPσ-deficient brains. (**D**) Genetic depletion of PTPσ results in a decreased distribution of full length APP (APP FL) in lipid raft membrane domains. Flotillin-1, a raft-associated protein, is shown as loading control. (**E**) Results from 8 pairs of age- and gender-matched mice, each with 1 wild type and 1 PTPσ-deficient, were quantified by densitometry. **B** and **E**, Data are mean ± SEM. *p* value, Student’s *t* test, 2-tailed. **C**, Bars represent means. *p* value and *R^2^*, linear regression.

## PTPσ modulates traffic signaling and APP lipid raft distribution

As previous studies substantiated an important role of membrane sorting in APP accessibility to the secretases, we followed on to analyze APP membrane distribution and pertinent traffic signaling in relation to PTPσ.

Lipid rafts, membrane microdomains enriched in sphingolipid and cholesterol, are critical subcellular locale for APP interaction with the β- and γ-secretases that is required for subsequent amyloidogenic processing (*25, 26*). The regulation of APP lipid raft trafficking, as established earlier, is mediated by Src kinase and its substrate, the Low-density Lipoprotein Receptor-related Protein 1 (LRP1). Via its cytoplasmic domain, LRP1 forms molecular complex with APP, which not only facilitates APP trafficking to the lipid rafts but also promotes BACE1-APP interaction and therefore APP β-processing (*27, 28*). These functions of LRP1 are dictated by the phosphorylation of its C-terminal tyrosine4507 (Y4507) (*28–30*), which in turn is enabled by Src kinase (*31*), another singling component that was shown crucial in regulating APP β-processing (*32, 33*).

In wild type mouse brains under physiological conditions, only a small fraction of APP is localized in the lipid raft membrane domains (fig. S3). In the absence of PTPσ, the level of raft-bound APP is further reduced (Fig. 4D, E). No alteration of BACE1 distribution, however, was observed in response to PTPσ depletion (data not shown). In support of this observation, analysis on APP traffic regulators showed consistent signaling (fig. S4). Genetic depletion of PTPσ results in deactivation of Src kinase, as indicated by increased phosphorylation of Src tyrosine527 (Y527) (fig. S4A, B), a signaling event that renders the kinase in an inactive folding. Src phospho-Y527 was previously reported as a substrate of PTPδ, another member of PTPσ phosphatase family (*34*). The enhanced Src phospho-Y527 level in the absence of PTPσ suggests that Src is also a substrate of PTPσ. In addition, presumably as a consequence of Src deactivation, we observed a reduction of LRP1 Y4507 phosphorylation exclusively within the raft fraction (fig. S4E, F), which, taking together with its role in APP raft-targeting and BACE1-APP interaction, may provide explanation for the decrease of raft-bound APP and BACE1-APP affinity in PTPσ-deficient brains. Collectively, these concerted findings suggest a possible PTPσ-Src-LRP1 signaling cascade in governing APP metabolism.

## APP metabolic regulation is dependent on PTPσ phosphatase activity

To further examine the role of PTPσ phosphatase activity, we tested a recombinant peptide, designed to bind PTPσ wedge domain and inhibit the phosphatase, in a litany of assays to assess APP β-processing. This peptide, known as PN001, is composed of 24 amino acids derived from PTPσ wedge domain along with a TAT sequence that facilitates membrane penetration (*35*). Our previous studies showed that PTPσ modulation by PN001 successfully overturns the signaling of chondroitin sulfate proteoglycans (CSPGs), a class of ligands that activate PTPσ activity (*11, 35, 36*). In pilot *in vitro* assays (using crude brain homogenates that contain partially intact cellular membranes, fig. S5A) and *ex vivo* assays (with freshly isolated mouse hippocampal tissues, fig. S5B), PN001 decreased APP CTFβ level, in comparison to the vehicle solution and a scrambled version of the peptide. More extensively, we analyzed the inhibitory effects of PN001 *in vivo*, via nasal administration to the wild type and TgAPP mice. The membrane permeable TAT sequence allows the recombinant peptide to access the brain, where PN001 manifests its inhibitory effects on the levels of CTFβ originated from both mouse endogenous APP (in the wild type mice, Fig. 5A, C) and human transgenic APP (in TgAPP-SwDI mice, Fig. 5B, D). Such effects are consistently reproduced in TgAPP-SwInd mice, but not observed upon PTPσ depletion (data not shown). In addition, when compared side-by-side within experiments, it is noticeable that PN001 showed a more congruent inhibitory performance than a BACE1 inhibitor Lanabecestat, that was previously in phase 3 Alzheimer clinical trial (Fig. 5D). Taken together, these data from various assays provided concordant evidence that APP metabolism is dependent on PTPσ phosphatase activity.

**Figure 5.**
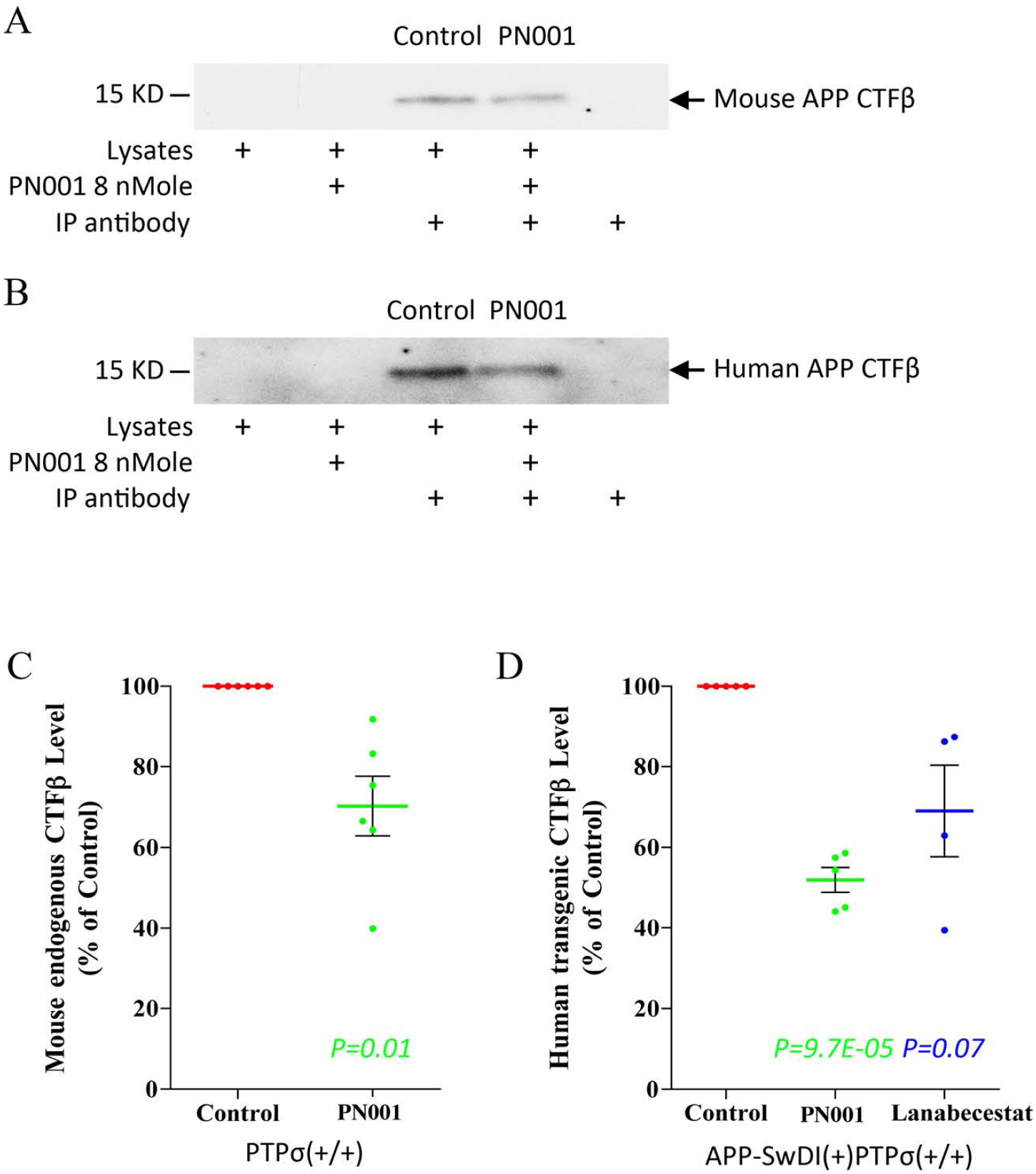
APP CTFβ level in the brain is dependent on PTPσ phosphatase activity. The membrane permeable TAT sequence in PN001 allows the recombinant peptide to access the brain and result in reduced levels of CTFβ originated from both mouse endogenous APP and human transgenic APP. Balb/c wild type mice (**A**) and TgAPP-SwDI mice (**B**) were treated via nasal administration of either vehicle solution (control) or PN001 (8 nmol). Forebrains were collected 30 minutes later and CTFβ levels were analyzed as in Fig.2 C and D. Mock immunoprecipitaion (IP) without antibody or lysates were used as negative controls. Results were quantified by densitometry. (**C**) Values from 6 repeated experiments as shown in **a**, each using a pair of gender-matched Balb/c wild type littermates. (**D**) Data plot from 5 repeated experiments as shown in **B**, using gender-matched TgAPP-SwDI littermates, among which 4 experiments included an additional mouse treated with BACE1 inhibitor Lanabecestat (also 8 nmol and nasal administration). Vehicle solution for each mouse contains 5 nmol of γ-secretase inhibitor DAPT to prevent CTFβ degradation. All values of treated samples were normalized to that of the control sample within each experiment. The *in vivo* efficacy of PN001 is consistent with results from *in vitro* and *ex vivo* assays (Fig. S5), suggesting a dependency of APP metabolism on PTPσ phosphatase activity. Line, mean. Error bar, SEM. *p* value, Student’s *t* test, 2-tailed, all compared to control.

Moreover, treatment with PN001 results in similar effects on the phosphorylation of Src Y527 and LRP1 Y4507 as observed in PTPσ-deficient brains (fig. S4C, D, G, H), further indicating a PTPσ-Src-LRP1 consecutive enzyme-substrate signaling cascade. In line with previous findings that APP β-processing is suppressed upon Src downregulation by either RNAi or PP2 inhibition (*32*), Src inhibitors (SKI-606 and WH-4-023) in our assays also led to a reduction of CTFβ level (data not shown) to a similar extent as PN001, providing additional support for the involvement of PTPσ downstream pathway in regulating APP processing.

## PTPσ depletion ameliorates AD-related neuroinflammation and synaptic impairment

It has been well established that overproduction of Aβ in the brain elicits multiplex downstream pathological events, including chronic inflammatory responses of the glia, such as persistent astrogliosis. The reactive (inflammatory) glia would then crosstalk with neurons, evoking a vicious feedback loop that exacerbates neurodegeneration during disease progression (*37–39*).

Among many existing AD mouse models, TgAPP-SwDI is one of the earliest to develop neurodegenerative pathologies and behavioral deficits (*17*). We therefore chose the model to further examine the role of PTPσ in downstream AD-related pathologies.

By the age of 9 months, severe neuroinflammation is developed in the brains APP-SwDI(+)PTPσ(+/+) mice, as measured by the level of GFAP (Glial Fibrillary Acidic Protein), a marker of astrogliosis (Fig. 6A-D; fig. S6). In the DG hilus, for example, GFAP expression level in the APP-SwDI(+)PTPσ(+/+) mice is more than tenfold compared to that in gender-matched non-transgenic littermates [APP-SwDI(-)PTPσ(+/+)]. PTPσ deficiency, however, effectively attenuates astrogliosis induced by the amyloidogenic transgene. In the APP-SwDI(+)PTPσ(−/−) brains, depletion of PTPσ restores GFAP expression in DG hilus back to a level close to that of non-transgenic wild type littermates (Fig. 6D).

**Figure 6.**
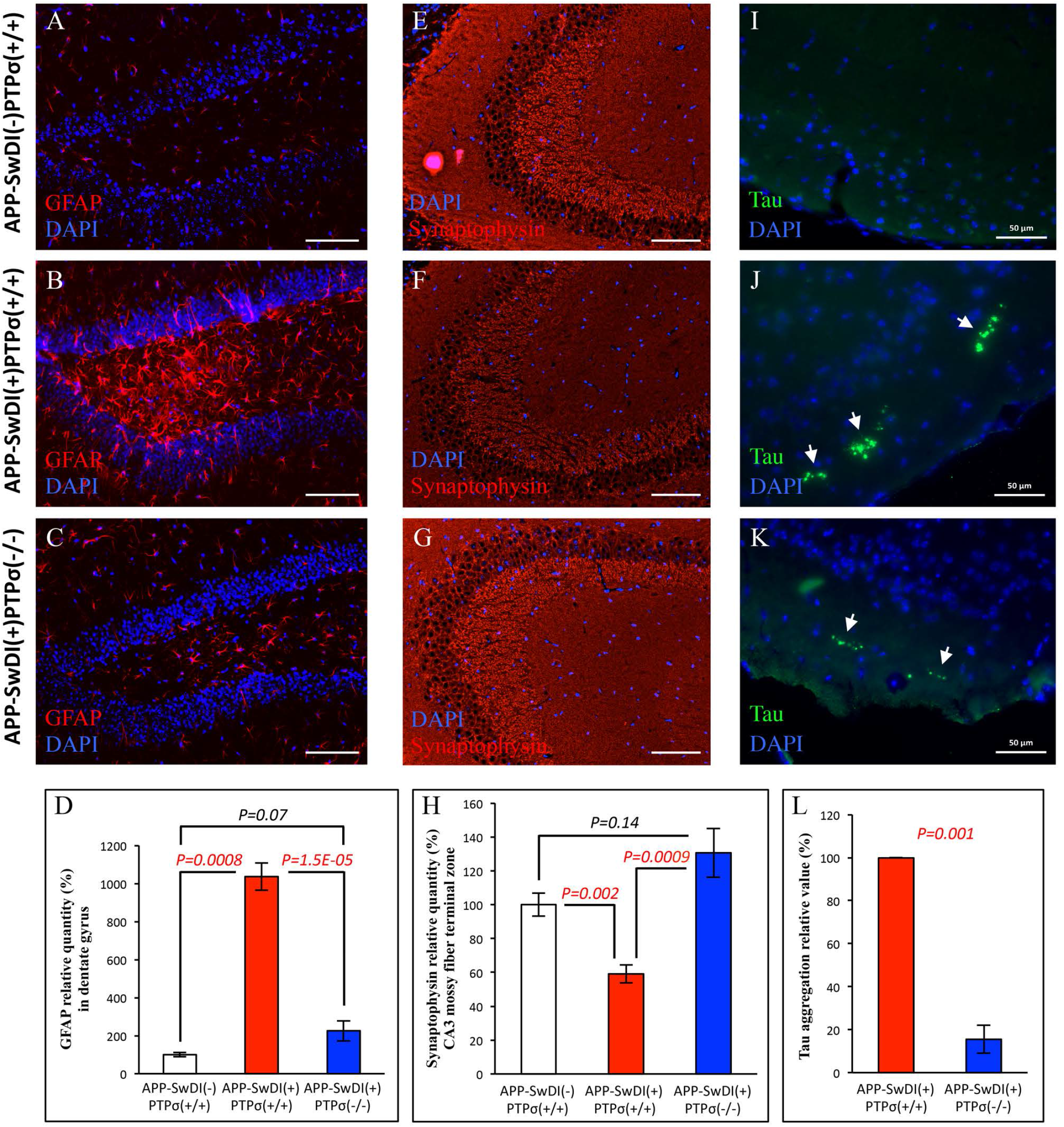
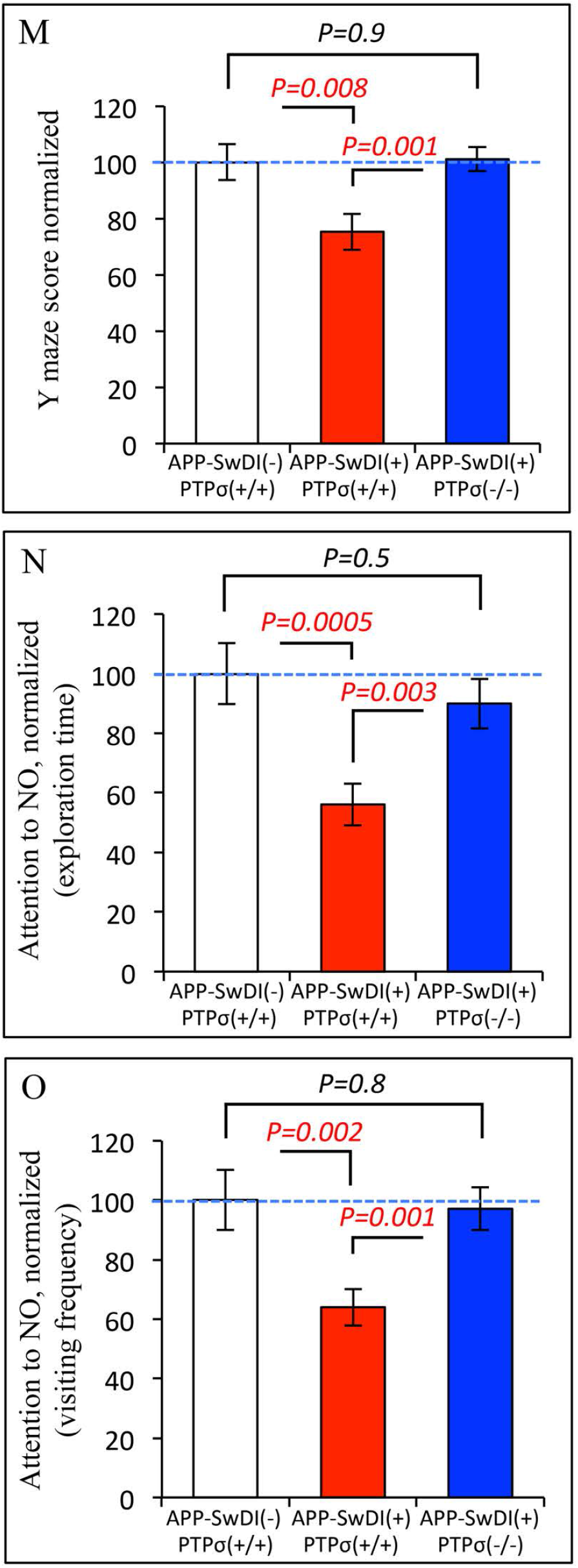
PTPσ PTPσ deficiency ameliorates AD-related pathologies and behavioral deficits. (**A**-**D**) PTPσ deficiency attenuates reactive astrogliosis in APP transgenic mice. Representative images show GFAP (red, a marker of reactive astrocytes) and DAPI staining of nuclei (blue) in the dentate gyrus (DG) of 9-month old TgAPP-SwDI mice with or without PTPσ, along with their non-transgenic wild type littermates. Images of GFAP in primary somatosensory cortex can be found in Fig S6. (**D**) ImageJ quantification of GFAP level in DG hilus from mice aged between 9 to 11 months. APP-SwDI(-)PTPσ(+/+), n=4; APP-SwDI(+)PTPσ(+/+), n=4; APP-SwDI(+)PTPσ(−/−), n=6. (**E**-**H**) PTPσ deficiency protects APP transgenic mice from synaptic loss. Representative images show immunofluorescent staining of presynaptic marker Synaptophysin in the mossy fiber terminal zone of CA3 region. Synaptophysin, red; DAPI, blue. Larger images can be found in Fig. S7. (**H**) ImageJ quantification of Synaptophysin expression level in CA3 mossy fiber terminal zone from mice aged between 9 to 11 months. APP-SwDI(-)PTPσ(+/+), n=4; APP-SwDI(+)PTPσ(+/+), n=6; APP-SwDI(+)PTPσ(−/−), n=6. (**I**-**L**) PTPσ deficiency diminishes Tau aggregation in APP transgenic mice. (**I**) No Tau aggregates are detected in non-transgenic wild type brains. (**J**) Representative images of many areas with Tau aggregation in the brains of gender-matched APP-SwDI(+)PTPσ(+/+) littermates. (**K**) Representative images of a few areas with Tau aggregation in gender-matched APP-SwDI(+)PTPσ(−/−) littermates. Tau, green; DAPI, blue. Arrows points to Tau aggregates. Images of Tau aggregates in the hippocampal region can be found in Fig. S8. (**L**) Bar graph shows quantification of Tau aggregation in coronal brain sections from 4 pairs of gender-matched APP-SwDI(+)PTPσ(+/+) and APP-SwDI(+)PTPσ(−/−) littermates of 9 to 11 month-old. Scale bars, 100 µm in **A-C** and **E-G**, 50 µm in **I-K**. For **D** and **H**, total integrated density of GFAP in DG hilus and Synaptophysin in CA3 mossy fiber terminal zone, respectively, was normalized to the area size to yield average intensity as shown in the bar graphs. Mean value of each group was normalized to that of non-transgenic APP-SwDI(-)PTPσ(+/+) mice. For **L**, values represent total integrated density from the entire coronal sections, and for each pair the value from APP-SwDI(+)PTPσ(−/−) sample is normalized to the value from APP-SwDI(+)PTPσ(+/+) sample. (**M**-**O**) PTPσ deficiency rescues behavioral deficits in APP transgenic mice. (**M**) In the Y-maze assay, performance of spatial navigation is scored by the percentage of spontaneous alternations among total arm entries. Compared to non-transgenic wild type mice, APP-SwDI(+)PTPσ(+/+) mice show deficit of short-term spatial memory, which is rescued by genetic depletion of PTPσ in APP-SwDI(+)PTPσ(−/−) mice. APP-SwDI(-)PTPσ(+/+), n=23 (18 females and 5 males); APP-SwDI(+)PTPσ(+/+), n=52 (30 females and 22 males); APP-SwDI(+)PTPσ(−/−), n=35 (22 females and 13 males). (**N**, **O**) Novel object test. NO, novel object. FO, familiar object. Attention to NO is measured by the ratio of NO exploration to total object exploration (NO+FO) in terms of exploration time (**N**) and visiting frequency (**O**). APP-SwDI(+)PTPσ(+/+) mice showed decreased interest in NO compared to wild type APP-SwDI(-)PTPσ(+/+) mice. The deficit is reversed by PTPσ depletion in APP-SwDI(+)PTPσ(−/−) mice. APP-SwDI(-)PTPσ(+/+), n=28 (19 females and 9 males); APP-SwDI(+)PTPσ(+/+), n=46 (32 females and 14 males); APP-SwDI(+)PTPσ(−/−), n=29 (21 females and 8 males). For both behavioral assays, ages of all groups are similarly distributed between 4 and 11 months. Values are normalized to that of non-transgenic APP-SwDI(-)PTPσ(+/+) mice within the colony. Raw data before normalization and additional analyses of the novel object test can be found in Fig. S12 and S13. All *p* values, Student’s *t* test, 2-tailed. All data are mean ± SEM.

Among all brain regions, the most affected by the expression of TgAPP-SwDI transgene appears to be the hilus of the DG, where Aβ deposition and astrogliosis are both found to be the most severe (Fig. 2G; Fig. 6A-D). We therefore questioned whether the pathologies in this area have an impact on the mossy fiber axons of DG pyramidal neurons, which project through the hilus into the CA3 region, where they synapse with the CA3 dendrites. Upon examining the presynaptic markers in CA3 mossy fiber terminal zone, we found decreased levels of Synaptophysin and Synapsin-1 in the APP-SwDI(+)PTPσ(+/+) mice, comparing to their gender-matched non-transgenic littermates (Fig. 6E-H; fig. S7, data not shown for Synapsin-1). Such synaptic impairment, evidently resulting from the expression of the APP transgene and probably the overproduction of Aβ, is reversed by genetic depletion of PTPσ in the APP-SwDI(+)PTPσ(−/−) mice (Fig. 6H).

Interestingly, we noticed that the APP-SwDI(+)PTPσ(−/−) mice sometimes express higher levels of presynaptic markers in the CA3 terminal zone than their non-transgenic wild type littermates (Fig. 6H). This observation, although not statistically significant in our quantification analysis, may suggest an additional synaptic effect of PTPσ that is independent of the APP transgene, as observed in previous studies (*40*).

## Tau pathology in aging AD mouse brains is dependent on PTPσ

Neurofibrillary tangles composed of hyperphosphorylated and aggregated Tau are commonly found in AD brains. These tangles tend to develop in a hierarchical pattern, appearing first in the entorhinal cortex before spreading to other brain regions (*6, 7*). The precise mechanism of tangle formation, however, is poorly understood. The fact that Tau tangles and Aβ deposits can be found in separate locations in postmortem brains has led to the question of whether Tau pathology in AD is independent of Aβ (*6, 7*). Additionally, quintessential neurofibrillary Tau tangles have not been reported in any of the APP transgenic mouse models, even in those with severe cerebral β-amyloidosis, further questioning the relationship between Aβ and Tau pathologies *in vivo*.

Nonetheless, a few studies did show non-tangle like assemblies of Tau within dystrophic neurites surrounding Aβ plaques in some APP transgenic mice (*41–43*), arguing that Aβ can be a causal factor for Tau dysregulation. In histological analysis using an antibody against the proline-rich domain of Tau, we observed diminutive aggregations in the brains of TgAPP-SwDI and TgAPP-SwInd mice during the course of aging (around 9 months for the APP-SwDI(+)PTPσ(+/+) mice and 15 months for the APP-SwInd(+)PTPσ(+/+) mice, but not in younger ages) (Fig. 6I-L; fig. S8 and 9). The sizes of the Tau aggregates are small to the extent that they may not be noticeable under lower magnification even with a signal amplification using a tertiary antibody, which could explain the lack of similar reports in the past. Such aggregations are not seen in gender-matched non-transgenic littermates (Fig. 6I), suggesting that it is a pathological event downstream from the expression of amyloidogenic APP transgenes and likely a result of Aβ cytotoxicity. Genetic depletion of PTPσ, which diminishes Aβ levels, suppresses Tau aggregation in both TgAPP-SwDI and TgAPP-SwInd mice (Fig. 6L; fig. S8 and 9).

In both TgAPP models, these small Tau aggregates are found predominantly in the molecular layer of the piriform and entorhinal cortices, and occasionally in the hippocampal region (fig. S8A), reminiscent of the early stage tangle locations in AD brains (*44*). Upon closer examination, the aggregates are not in typical tangle morphologies, but often in punctate shapes, likely in debris of degenerated cell bodies and neurites, scattered in areas mostly free of nuclear staining (fig. S10A-F). Rarely, a few are in fibrillary structures, probably in degenerated cells before disassembling (fig. S10G, H). Because such Tau pathology in TgAPP mice is an unprecedented finding, we reconfirmed the results using an additional antibody against the C-terminus of Tau (fig. S10I). Both antibodies detected the same morphologies and distribution pattern of the Tau aggregates.

Consistent with the findings in postmortem AD brains, the presence of Tau aggregates in the TgAPP mouse brains does not correlate with Aβ deposition. At the age of 9 months, Aβ deposition in TgAPP-SwDI brains is pronounced in the hippocampus yet only sporadic in the piriform and entorhinal cortex where most Tau aggregates are found (Fig. 2G, H; fig. S8 and 9). Given that the causation of Tau pathology in these mice is probably related to the overproduced Aβ, the segregation of predominant areas for Aβ and Tau depositions may indicate that the cytotoxicity originates from soluble Aβ rather than the deposited amyloid. It is also evident that neurons in different brain regions are not equally vulnerable to developing Tau pathology.

We next examined whether the expression of APP transgenes or genetic depletion of PTPσ regulates Tau aggregation by changing its expression level and/or phosphorylation status. Western blot analysis of brain homogenates showed that Tau protein expression is not affected by the APP transgenes or PTPσ (fig. S11), suggesting that the aggregation may result from local misfolding of Tau rather than an overexpression of the protein. Additionally, both biochemical analyses with brain homogenates and immunohistological staining of the Tau aggregates revealed that the TgAPP transgenes, which apparently cause Tau aggregation, do not enhance the phosphorylation of Tau residues including Serine191, Therionine194, and Therionine220 (data not shown), whose homologues in human Tau (Serine202, Therionine205, and Therionine231) are typically hyperphosphorylated in neurofibrillary tangles. Consistent with our findings, a recent quantitative study showed that post-translational modifications of Tau in the TgAPP-SwInd mice are similar to that in wild type mice (*45*).

Although the underlying mechanism that triggers Tau misfolding is still unclear, the finding of Tau pathology in these mice establishes a causal link between the expression of amyloidogenic APP transgenes and a dysregulation of Tau assembly. The mitigated Tau aggregation upon PTPσ depletion may due to the reduction of amyloidogenic metabolites of APP.

Malfunction of Tau is broadly recognized as a neurodegenerative marker as it indicates microtubule deterioration (*8*). The ameliorating effect on Tau aggregation by PTPσ deficiency thus provides additional evidence for a pivotal role of this cell surface receptor in regulating neuronal integrity.

## PTPσ deficiency rescues AD-related behavioral deficits

We next assessed whether the alleviation of neuropathologies by PTPσ depletion is accompanied with a rescue from AD-relevant behavioral deficits. The most common symptoms of AD include short-term memory loss and apathy among the earliest, followed by spatial disorientation amid impairment of many other cognitive functions as the dementia progresses. Using Y maze and novel object assays as surrogate models, we evaluated these cognitive and psychiatric features in the TgAPP-SwDI and TgAPP-SwInd mice.

The Y-maze assay, which allows mice to freely explore three identical arms, measures their short-term spatial memory. It is based on the natural tendency of mice to alternate arm exploration without repetitions. The performance is scored by the percentage of spontaneous alternations among total arm entries, and a higher score indicates better spatial navigation. Compared to the non-transgenic wild type mice within the colony, the APP-SwDI(+)PTPσ(+/+) mice show a clear deficit in their performance. Genetic depletion of PTPσ in the APP-SwDI(+)PTPσ(−/−) mice, however, unequivocally restores the cognitive performance back to the level of non-transgenic wild type mice (Fig. 6M; fig. S12).

Apathy, the most common neuropsychiatric symptom reported among individuals with AD, is characterized by a loss of motivation and diminished attention to novelty, and has been increasingly adopted into early diagnosis of preclinical and early prodromal AD (*46–49*). Many patients in early stage AD lose attention to novel aspects of their environment despite their ability to identify novel stimuli, suggesting an underlying defect in the circuitry responsible for further processing of the novel information (*47, 48*). As a key feature of apathy, such deficits in attention to novelty can be accessed by the “curiosity figures task” or the “oddball task” in patients (*47, 48, 50*). These visual-based novelty encoding tasks are very similar to the novel object assay for rodents, which measures the interest of animals in a novel object (NO) when they are exposed simultaneously to a pre-familiarized object (FO). We therefore used this assay to test the attention to novelty in the APP transgenic mice. When mice are pre-trained to recognize the FO, their attention to novelty is then measured by the discrimination index denoted as the ratio of NO exploration to total object exploration (NO+FO), or alternatively, by the ratio of NO exploration to FO exploration. Whereas both ratios are commonly used, a combination of these assessments provides a more comprehensive evaluation of animal behavior. In this test, as indicated by both measurements, the expression of APP-SwDI transgene in the APP-SwDI(+)PTPσ(+/+) mice leads to a substantial decrease in NO exploration as compared to non-transgenic wild type mice (Fig. 6N, O; fig. S13). Judging by their NO/FO ratios, it is evident that the transgenic and non-transgenic groups are both able to recognize and differentiate between the two objects (fig. S13A, B). Thus, the reduced NO exploration by the APP-SwDI(+)PTPσ(+/+) mice may reflect a lack of interest in the NO or an inability to shift attention to the NO. Once again, this behavioral deficit is largely reversed by PTPσ deficiency in the APP-SwDI(+)PTPσ(−/−) mice (Fig. 6N, O; fig. S13), consistent with previous observation of increased NO preference in the absence of PTPσ (*40*).

To further verify the effects of PTPσ on these behavioral aspects, we additionally tested the TgAPP-SwInd mice in both assays and observed similar results, confirming an improvement on both short-term spatial memory and attention to novelty upon genetic depletion of PTPσ (fig. S14).

## Discussion

Here we report that cerebral accumulation of β-amyloidogenic peptides and several downstream disease features are dependent on PTPσ in two mouse models of genetically inherited AD. This form of AD develops inevitably in people who carry gene mutations that promote amyloidogenic processing of APP. Our data suggest that targeting PTPσ is a potential therapeutic approach to overcome such dominant genetic driving forces and curtail AD progression. An advantage of this targeting strategy is that it suppresses the overproduction of APP amyloidogenic metabolites without broadly affecting other major substrates of the β- and γ-secretases, thus predicting a more promising translational potential as compared to those in clinical trials that broadly inhibit the secretases.

PTPσ was previously characterized as a neuronal receptor of the chondroitin sulfate- and heparan sulfate-proteoglycans (CSPGs and HSPGs) (*9, 10*). In response to these two classes of extracellular ligands, PTPσ functions as a “molecular switch” by regulating neuronal behavior in opposite manners (*11*). A pivotal role for this proteoglycan sensor PTPσ in AD-like pathogenesis may therefore implicate the involvement of perineuronal matrix in AD etiology.

More than 95% of AD cases are sporadic, which are not genetically inherited but likely result from insults to the brain that occurred earlier in life. AD risk factors, such as traumatic brain injury and cerebral ischemia (*51–54*), have been shown to induce overproduction of Aβ in both human and rodents (*55–59*), and speed up progression of this dementia in animal models (*60–62*). However, what promotes the amyloidogenic processing of APP in these situations is still a missing piece of the puzzle in understanding the AD-causing effects of these notorious risk factors.

Notably, both traumatic brain injury and cerebral ischemia result in pronounced remodeling of the perineuronal microenvironment at lesion sites, marked by increased expression of CSPGs (*63–66*), a major component of the perineuronal net that is upregulated during neuroinflammation and glial scar formation (*67–69*). In the brains of AD patients, CSPGs were found associated with Aβ depositions, further suggesting an uncanny involvement of these proteoglycans in AD development (*70*). On the other hand, analogues of heparan sulfate (HS, carbohydrate side chains of HSPGs that bind to PTPσ) were shown to inhibit BACE1 activity on APP, suggesting their function in preventing the initiation of “Aβ cascade” (*71*). Furthermore, the expression of Heparanase, an enzyme that degrades HS, was found markedly increased after cerebral ischemia and in AD brains (*72, 73*). Collectively, these findings revealed that disrupted molecular balance in the perineuronal environment by chronic upregulation of CSPGs or degradation of HSPGs is a common feature in brains of Alzheimer patients and those suffered from the risk factors.

We hence speculate that sustained aberrant signaling of the proteoglycans in the lesioned brains, readily mediated by neuronal receptor PTPσ, may lead to biased processing of APP and subsequently ignite insidious pathological cascades that precipitate the onset of AD. As such, restoring the integrity of brain microenvironment may be essential in preventing sporadic AD for the population at risk, and PTPσ, a gatekeeper of perineuronal signaling, would therefore provide a molecular target for therapeutic intervention.

Taken together with the role of PTPσ previously established in axonal regeneration, the recuperative efficacy of PTPσ downregulation, via genetic depletion or pharmacological inhibition, in both disease models of AD and CNS injury (*10, 35*), illustrates a common underlying mechanism that is mediated by the signaling of this neuronal receptor.

## Acknowledgements

We thank Dennis Selkoe, Jared Cregg, and Miriam Osterfield for scientific discussion; Anthony Brown, Christopher Edwards, Bradley Lang, Patricia Hess, Yan Wang for comments on the manuscript; Lennart Mucke for providing the TgAPP-SwInd (J20) mice; William E. Van Nostrand for generating the TgAPP-SwDI mice; Jonathan Cherry for technical support on imaging; Li Zhang for technical advice on rodent pathology. The behavioral data were collected with the support of National Institute of Neurological Disorders and Stroke Grant P30 NS045758. Aβ ELISA analyses were performed with the support of R01 AG043522-01 to Landreth (PI) by National Institutes of Health. The rest of the work was supported by Ohio State University Startup Funds and Sponsored Research Grant from Westfield Ventures LLC to Shen (PI).

## Author contributions

YG, YS, AWC, KX, AFY, SC, and ML conducted experiments and data analyses, under the supervision of GEL, RJN, and Y Shen. YG, KX, and AWC also provided comments on the manuscript. MLT provided PTPσ-deficient mice, advised on PTPσ characterization, and commented on the manuscript. JS, GEL, and RJN advised on experimental design and commented on the manuscript. Y Shen conceived the project, designed the study, and wrote the paper.

*Note: as there are 2 authors whose initials are YS, the last author is noted as“Y Shen”*.

## Supplementary Materials

### Materials and Methods

#### Mouse lines

Mice were maintained under standard conditions approved by the Institutional Animal Care and Use Committee. Wild type and PTPσ-deficient mice of Balb/c background were provided by Dr. Michel L. Tremblay (*74*). Homozygous TgAPP-SwDI mice, C57BL/6-Tg(Thy1-APPSwDutIowa)BWevn/Mmjax, stock number 007027, were from the Jackson Laboratory. These mice express human APP transgene harboring Swedish, Dutch, and Iowa mutations, and were bred with Balb/c mice heterozygous for the PTPσ gene knockout allele to generate bigenic mice heterozygous for both TgAPP-SwDI and PTPσ genes, which are hybrids of 50% C57BL/6J and 50% Balb/c genetic background. These mice were further bred with Balb/c mice heterozygous for the PTPσ gene and the offsprings were used in experiments, which include littermates of the following genotypes: TgAPP-SwDI(+/−)PTPσ(+/+), mice heterozygous for TgAPP-SwDI transgene with wild type PTPσ; TgAPP-SwDI(+/−)PTPσ(−/−), mice heterozygous for TgAPP-SwDI transgene with genetic depletion of PTPσ; TgAPP-SwDI(−/−)PTPσ(+/+), mice free of TgAPP-SwDI transgene with wild type PTPσ. Both TgAPP-SwDI(−/−)PTPσ(+/+) and Balb/c PTPσ(+/+) are wild type mice but with different genetic background. Heterozygous TgAPP-SwInd (J20) mice, 6.Cg-Tg(PDGFB-APPSwInd)20Lms/2Mmjax, were provided by Dr. Lennart Mucke. These mice express human APP transgene harboring Swedish and Indiana mutations, and were bred with the same strategy as described above to obtain mice with genotypes of TgAPP-SwInd (+/−)PTPσ(+/+) and TgAPP-SwInd (+/−)PTPσ(−/−).

For all experiments expect immunohistochemistry, animals were euthanized using CO_2_.

#### Antibodies

**Antibodies:**
| Primary Antibodies | Application | Clone # | Catalog # | Supplier |
| --- | --- | --- | --- | --- |
| Mouse anti-Actin | WB | AC-40 | A4700 | Sigma-Aldrich |
| Rabbit anti-APH1 | WB |  | PA5-20318 | Thermo Scientific |
| Rabbit anti-APP C-term | WB, IP, IHC | Y188 | NB110-55461 | Novus Biologicals |
| Mouse anti-murine Aβ, 1-16 | WB, IP | M3.2 | 805701 | Biolegend |
| Mouse anti-human Aβ, 1-16 | WB, IP, IHC, ELISA | 6E10 | 803016 | Biolegend |
| Mouse anti-Aβ, 17-24 | WB, IHC | 4G8 | SIG-39220 | Biolegend |
| Mouse HRP-conjugated anti-Aβ 1-40 | ELISA | 11A50-B10 | SIG-39146 | Biolegend |
| Mouse HRP-conjugated anti-Aβ 1-42 | ELISA | 12F4 | 805507 | Biolegend |
| Rabbit anti-BACE1 C-Term, B690 | WB |  | PRB-617C | Covance |
| Guinea Pig anti-BACE1 C-Term | IP |  | 840201 | Biolegend |
| Rabbit anti-Caveolin1 | WB |  | 3238 | Cell Signaling |
| Rabbit anti-Flotillin1 | WB |  | 3253 | Cell Signaling |
| Chicken anti-GFAP | IHC |  | ab4674 | Abcam |
| Rabbit anti-LRP1 C-term | WB |  | ab92544 | Abcam |
| Rabbit anti-LRP1 Phospho-Tyr4507 | WB |  | sc33049 | Santa Cruz Biotechnology |
| Rabbit anti-Neuregulin | WB |  | sc-348 | Santa Cruz Biotechnology |
| Rabbit anti-Nicastrin | WB |  | 5665 | Cell Signaling |
| Rabbit anti-Notch NICD (val1744) | WB |  | 4147 | Cell Signaling |
| Rabbit anti-Notch (C-20) | WB |  | sc-6014R | Santa Cruz Biotechnology |
| Rabbit anti-PEN2 | WB |  | 8598 | Cell Signaling |
| Rabbit anti-Presenilin 1/2 NTF | WB |  | 840201 | Abcam |
| Rabbit anti-Presenilin 1 CTF | WB |  | 5643 | Cell Signaling |
| Rabbit anti-Presenilin 2 CTF | WB |  | 9979 | Cell Signaling |
| Mouse anti-PTPσ ICD | WB, IHC | 17G7.2 | MM-002-P | Medimabs |
| Mouse anti-PTPσ ECD | WB | | ab55640 | Abcam |
| Mouse anti-Src | WB |  | SM2591 | ECM Biosciences |
| Mouse anti-Src Phospho-Tyr527 | WB |  | SM2611 | ECM Biosciences |
| Rabbit anti-Synaptophysin | IHC |  | AB9272 | Millipore |
| Mouse anti-Tau | WB, IHC | Tau-5 | MAB361 | Millipore |
| Mouse anti-Tau | IHC | Tau-46 | 4019 | Cell Signaling |
| Secondary and Tertiary Antibodies | Application | Clone # | Catalog # | Supplier |
| Goat anti-mouse IgG HRP-conjugated | WB |  | 7076S | Cell Signaling |
| Goat anti-rabbit IgG HRP-conjugated | WB |  | 7074S | Cell Signaling |
| Goat anti-mouse IgG Alexa488 | IHC |  | A-11001 | Invitrogen |
| Donkey anti-goat IgG Alexa488 | IHC |  | A-11055 | Invitrogen |
| Chicken anti-rabbit IgG CF568 | IHC |  | SAB4600426 | Sigma-Aldrich |
| Donkey anti-chicken IgG Cy3 | IHC |  | 703-165-155 | JacksonImmunoResearch |

#### Immunohistochemistry

Adult rat and mice were perfused intracardially with fresh made 4% paraformaldehyde in cold phosphate-buffered saline (PBS). The brains were collected and post-fixed for 2 days at 4 °C. Paraffin embedded sections of 10 µM thickness were collected for immunostaining. The sections were deparaffinized and sequentially rehydrated. Antigen retrieval was performed at 100 °C in Tris-EDTA buffer (pH 9.0) for 50 minutes. Sections were subsequently washed with distilled water and PBS, incubated at room temperature for 1 hour in blocking buffer (PBS, with 5% normal donkey serum, 5% normal goat serum, and 0.2% Triton X-100). Primary antibody incubation was performed in a humidified chamber at 4°C overnight. After 3 washes in PBS with 0.2% Triton X-100, the sections were then incubated with a mixture of secondary and tertiary antibodies at room temperature for 2 hours. All antibodies were diluted in blocking buffer with concentrations recommended by the manufacturers. Mouse primary antibodies were detected by goat anti-mouse Alexa488 together with donkey anti-goat Alexa488 antibodies; rabbit primary antibodies were detected by chicken anti-rabbit CF568 and donkey anti-chicken Cy3 antibodies; chicken antibody was detected with donkey anti-chicken Cy3 antibody. Sections stained with only secondary and tertiary antibodies (without primary antibodies) were used as negative controls. At last, DAPI (Invitrogen, 300 nM) was applied on sections for nuclear staining. Sections were washed 5 times before mounted in Fluoromount (SouthernBiotech, Catalog # 0100-01).

Wide field and confocal images were captured using Zeiss Axio Imager M2 and LSM780, respectively. Images are quantified using the Zen 2 Pro software and ImageJ.

#### Protein extraction, immunoprecipitation, and western blot analysis

For the co-immunoprecipitation of APP and PTPσ, RIPA buffer was used (50 mM Tris-HCl, pH 8.0, 1 mM EDTA, 150 mM NaCl, 1% NP40, 0.1% SDS, 0.5% sodium deoxycholate). For the co-immunoprecipitation of APP and BACE1, NP40 buffer was used (50 mM Tris-HCl, pH 8.0, 1 mM EDTA, 150 mM NaCl, 1% NP40) without or with SDS at concentration of 0.1%, 0.3%, and 0.4%. For total protein extraction and immunopurification of CTFβ, SDS concentration in RIPA buffer was adjusted to 1% to ensure protein extraction from the lipid rafts. Mouse or rat forebrains were homogenized thoroughly on ice in designated homogenization buffer containing protease and phosphatase inhibitors (Thermo Scientific, Catalog # A32959). For each half of forebrain, buffer volume of at least 5 ml for mouse and 8 ml for rat was used to ensure sufficient detergent/tissue ratio. The homogenates were incubated at 4°C for 1 hour with gentle mixing, sonicated on ice for 2 minutes in a sonic dismembrator (Fisher Scientific Model 120, with pulses of 50% output, 1 second on and 1 second off), followed with another hour of gentle mixing at 4°C. All samples were used fresh without freeze-thaw to avoid artifacts due to protein precipitation.

For co-immunoprecipitation, the homogenates were then centrifuged at 85,000 x g for 1 hour at 4°C and the supernatants were collected. Protein concentration was measured using BCA Protein Assay Kit (Thermo Scientific, Catalog # 23227). 0.5 mg total proteins of brain homogenates were incubated with 5 μg of designated antibody and 30 μl of Protein-A agarose beads (50% slurry, Roche catalog # 11243233001), in a total volume of 1 ml adjusted with homogenization buffer. CTFβ immunopurification was carried out with the same procedure except that sample volumes were adjusted to 1 ml using RIPA buffer so that the final concentration of SDS was no more than 0.2%. Samples were gently mixed at 4°C overnight. Subsequently, the beads were washed 5 times with cold immunoprecipitation buffer. Samples were then incubated in Laemmli buffer with 100 mM of DTT at 75°C for 20 minutes and subjected to western blot analysis.

For analysis of protein expression level, the homogenates were centrifuged at 23,000 x g for 30 min at 4°C and the supernatants were collected. Protein concentration was measured using BCA Protein Assay Kit. For western blot analysis, sample loading was normalized to equal total protein amount per lane, ranging from 5 to 30 μg.

Electrophoresis of protein samples was conducted using 4-12% or 12% Bolt Bis-Tris Plus Gels, with either MOPS or MES buffer and Novex Sharp Pre-stained Protein Standard (Invitrogen catalog # LC5800). Proteins were transferred to nitrocellulose membrane (0.2 μm pore size, Bio-Rad) and blotted with selected antibodies (see table above) at concentrations recommended by the manufacturers. Primary antibodies were diluted in SuperBlock TBS Blocking Buffer (Thermo Scientific catalog # 37535) and incubated with the nitrocellulose membranes at 4°C overnight; secondary antibodies were diluted in PBS with 5% nonfat milk and 0.2% Tween20 and incubated at room temperature for 2 hours. Membranes were washes 4 times in PBS with 0.2% Tween20 between primary and secondary antibodies and before chemiluminescent detection with SuperSignal West Pico Chemiluminescent Substrate (Thermo Scientific, catalog # 34080).

Western blot band intensity was quantified by densitometry.

#### Aβ ELISA assays

Mouse forebrains were thoroughly homogenized in tissue homogenization buffer (2 mM Tris pH 7.4, 250 mM sucrose, 0.5 mM EDTA, 0.5 mM EGTA) containing protease inhibitor cocktail (Roche, catalog # 11697498001), followed by centrifugation at 135,000 x g (33,500 RPM with SW50.1 rotor) for 1 hour at 4°C. Proteins in the pellets were extracted with formic acid (FA) and centrifuged at 109,000 x g (30,100 RPM with SW50.1 rotor) for 1 hour at 4°C. The supernatants were collected and diluted 1:20 in neutralization buffer (1 M Tris base, 0.5 M Na_2_HPO_4_, 0.05% NaN_3_) and subsequently 1:3 in ELISA buffer (PBS with 0.05% Tween-20, 1% BSA, and 1 mM AEBSF). Diluted samples were loaded onto ELISA plates (Thermo Scientific, catalog # 469949) pre-coated with 6E10 antibody (see table above) to capture Aβ peptides. Serial dilutions of synthesized human Aβ 1-40 or 1-42 (American Peptide, catalog # 62-0-78, 62-0-80) were loaded to determine a standard curve. Aβ was detected using an HRP labeled antibody for either Aβ 1-40 or 1-42 (see table above). ELISA was developed using TMB substrate (Thermo Scientific, catalog # 34021) and reaction was stopped with 1N HCl. Plates were read at 450nm and concentrations of Aβ in samples were determined using the standard curve.

#### Non-raft and lipid raft membrane fractionation

Subcellular fractions from brain homogenates were separated based on their detergent solubility. The detergent soluble and resistant fractions are used as surrogates for non-raft and lipid raft membrane fractions, respectively. Changes in protein levels and signaling associated with the detergent resistant fractions are likely derived from native raft domains. Freshly isolated mouse brains were processed in serial steps of protein extraction, using buffers provided in subcellular protein fractionation kit (Thermo Scientific, catalog # 87790) with modified protocol. Tissues were homogenated in ice-cold MEB buffer provided in the kit, or using an equivalent NP40 buffer (50 mM Tris-HCl, pH 8.0, 1 mM EDTA, 150 mM NaCl, 1% NP40), followed with incubation on ice for 10 minutes. The homogenates were then centrifuged at 3000 × g for 5 minutes at 4°C and the supernatants collected were used as the “non-raft” fraction, which contains proteins soluble in mild non-ionic detergent, including most native non-raft membrane proteins, cytosolic proteins, and some raft proteins associated to the lipid rafts with low affinity. The (detergent resistant) pellets were further processed as described in the protocol provided with the kit, which include incubation with high salt concentration at room temperature and 37°C with nuclease to collect nuclear proteins and DNA-bound proteins respectively. The resulting pellets were then dissolved in PEB buffer (or alternatively RIPA buffer with 1%SDS) by incubation at room temperature for 10 minutes followed by centrifugation at 16,000 x g for 5 minutes. The final supernatants collected were used as the “raft fraction”, since this fraction contains lipid raft membrane proteins insoluble to mild detergent and high salt extraction, as verified by the enrichment of Caveolin1. For analysis of protein phosphorylation, the procedures may be simplified into two steps on ice to avoid signal attenuation at higher temperature. Following MEB extraction to obtain the non-raft fraction, the pellets were directly suspended in ice-cold RIPA buffer with 1% SDS, sonicated on ice for 2 minutes and gently mixed for 1 hour at 4°C. After centrifugation at 23,000 x g for 20 minutes, the supernatants were collected and used as the “lipid raft membrane fraction”. Protease and phosphatase inhibitors (Thermo Scientific, Catalog # A32959) were added to all buffers immediately before use.

#### APP β-processing assays

PN001, NH2-grkkrrqrrrcDMAEHMERLKANDSLKLSQEYESI-NH2; sPN001 (scrambled PN001), NH2-grkkrrqrrrcIREDDSLMLYALAQEKKESNMHES-NH2. Both peptides were synthesized by CSBio (>98% purity). The membrane permeable TAT sequence (in lower case) allows peptide penetration into the hippocampus tissue (in *ex vivo* assays) and the brain (in *in vivo* assays).

The *in vitro* assays were performed using detergent free brain homogenates from Balb/c wild type mice that contain partially intact cellular membranes. Freshly isolated mouse brains were homogenated on ice in HBSS buffer (Thermo Scientific, Catalog # 14025092) with 1 μM of γ-secretase inhibitor DAPT (Selleckchem, catalog # S2215) to prevent CTFβ degradation. The crude brain homogenates were aliquoted in equal volume and incubated at 37°C for 30 minutes with or without PN001 or sPN001, each at 5 μM final concentration. BACE1 function in the assay system was verified as CTFβ level gradually increases within the timeframe.

In *ex vivo* assays, the left and right hippocampal tissues were freshly isolated from each Balb/c wild type mouse brain, and randomly chosen to be incubated with or without 5 μM of PN001, at 37°C for 30 minutes with 1 μM of DAPT in artificial CSF (148 mM NaCl, 3 mM KCl, 1.4 mM CaCl_2_, 0.8 mM MgCl_2_, 0.8 mM Na_2_HPO_4_, 0.2 mM NaH_2_PO_4_).

In *in vivo* assays, gender-matched littermates were randomly selected to receive intranasal administration of either vehicle solution (10 μl, 5 nMole of DAPT and 7% DMSO in ddH_2_O), PN001 (8 nMole in 10 μl of vehicle solution), or BACE1 inhibitor Lanabecestat (also known as AZD3293 or LY3314814, Selleckchem, catalog # S8193, 8 nMole in 10 μl of vehicle solution). All reagents were prepared from stock solutions stored at −80°C and used fresh without repeated freeze-thaw. Stock solutions: PN001 at 4 mM in ddH_2_O; Lanabecestat at 40 mM in DMSO; DAPT at 10 mM in DMSO. Mice received 5 μl of solution in each nostril, and were held with nostrils facing up for 1 minute to allow absorption of the drugs before released back to cage. Immediate release of mice after nasal administration may result in less absorption and subpar efficacy as shown in the two highest data points of PN001 treatment in Fig. 5c. Forebrains were collected 30 minutes later.

Proteins were harvested for further analysis by the end of the assays. For *in vitro* assays, 2X RIPA buffer with 2% SDS was mixed with the homogenates at 1:1 ratio. For *ex vivo* and *in vivo* assays, the hippocampal tissues and forebrains were homogenated in RIPA buffer with 1% SDS. All lysis buffers contain protease and phosphatase inhibitors (Thermo Scientific, Catalog # A32959). All following steps for total protein extraction and immunopurification of APP CTFβ were carried out using the methods described above. Alternatively, forebrains from *in vivo* assays were processed to isolate subcellular fractions as described above.

#### Behavior assays

The Y-maze assay: Mice were placed in the center of the Y-maze and allowed to move freely through each arm. Their exploratory activities were recorded for 5 minutes. An arm entry is defined as when all four limbs are within the arm. For each mouse, the number of triads is counted as “spontaneous alternation”, which was then divided by the number of total arm entries, yielding a percentage score. The novel object test: On day 1, mice were exposed to empty cages (45 cm x 24 cm x 22 cm) with blackened walls to allow exploration and habituation to the arena. During day 2 to day 4, mice were returned to the same cage with two identical objects placed at an equal distance. On each day mice were returned to the cage at approximately the same time during the day and allowed to explore for 10 minutes. Cages and objects were cleaned with 70% ethanol between each animal. Subsequently, 2 hours after the familiarization session on day 4, mice were put back to the same cage where one of the familiar objects (randomly chosen) was replaced with a novel object, and allowed to explore for 5 minutes. Mice were scored using Observer software (Noldus) on their time duration and visiting frequency exploring either object. Object exploration was defined as facing the object and actively sniffing or touching the object, whereas any climbing behavior was not scored. The discrimination indexes reflecting interest in the novel object is denoted as either the ratio of novel object exploration to total object exploration (NO/(NO+FO)) or the ratio of novel object exploration to familiar object exploration (NO/FO). All tests and data extraction were conducted in a double-blinded manner.

#### Experimental replicates

The number of independent repeats (n) is indicated in the figure legends. All independent experiments were conducted with distinct samples.

#### Statistics

2-tailed Student’s *t* test was used for two-group comparison. Relationship between two variables (SDS concentration and APP-BACE1 association, as in Fig. 4c) was analyzed using linear regression. All data are mean values and error bars show standard error of the means (SEM).

---

**Figure S1.**
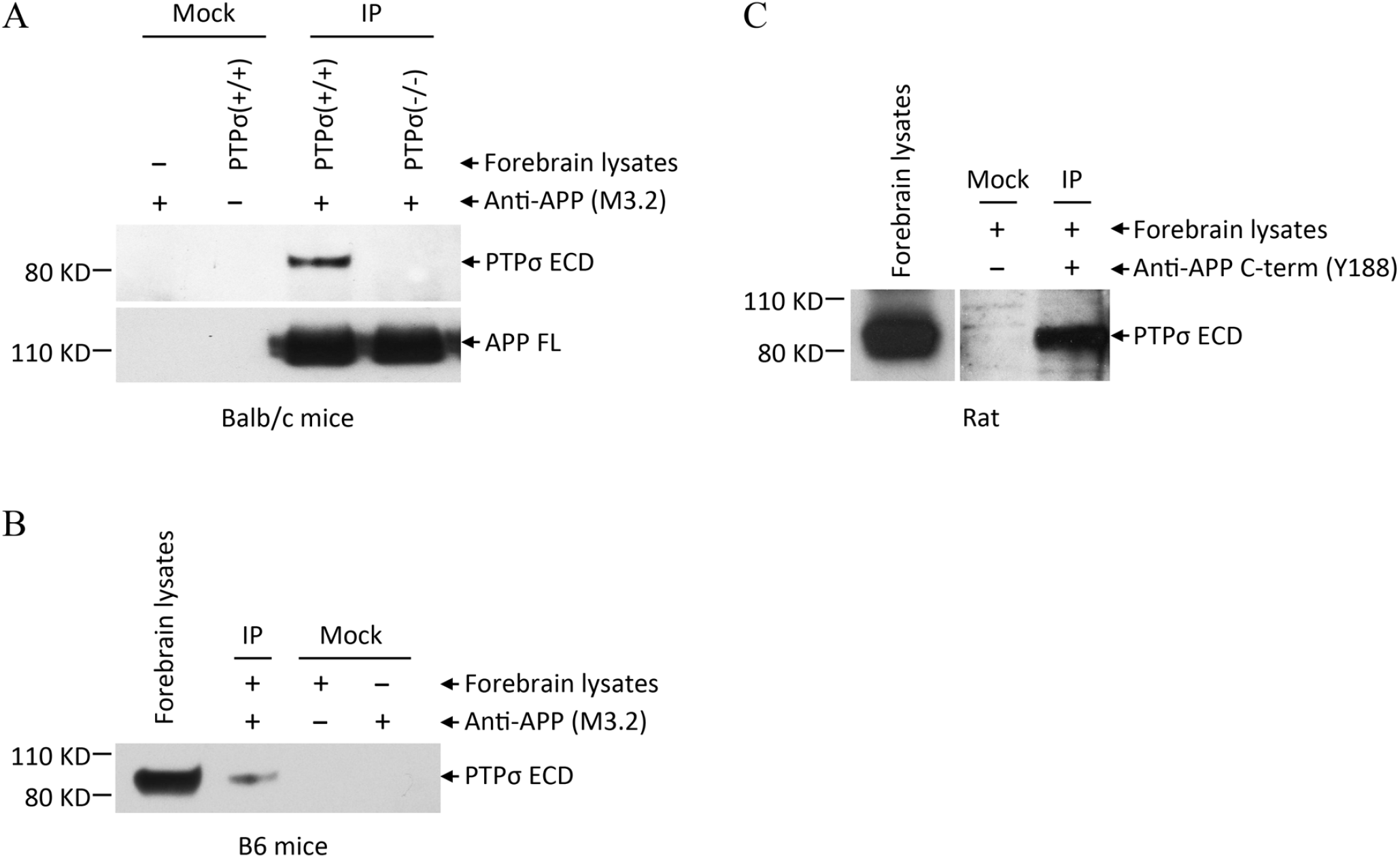
Molecular complex of PTPσ and APP in brains of various rodent species. (**A, B**) Co-immunoprecipitation using an anti-APP antibody specific for amino acid residues 1-16 of mouse Aβ (clone M3.2). PTPσ and APP binding interaction is detected in forebrains of Balb/c (**A**) and B6 (**B**) mice. (**C**) PTPσ co-immunoprecipitates with APP from rat forebrain lysates using an antibody (Y188) specific for the C-terminus of APP. Images shown are representatives of at least three independent experiments.

**Figure S2.**
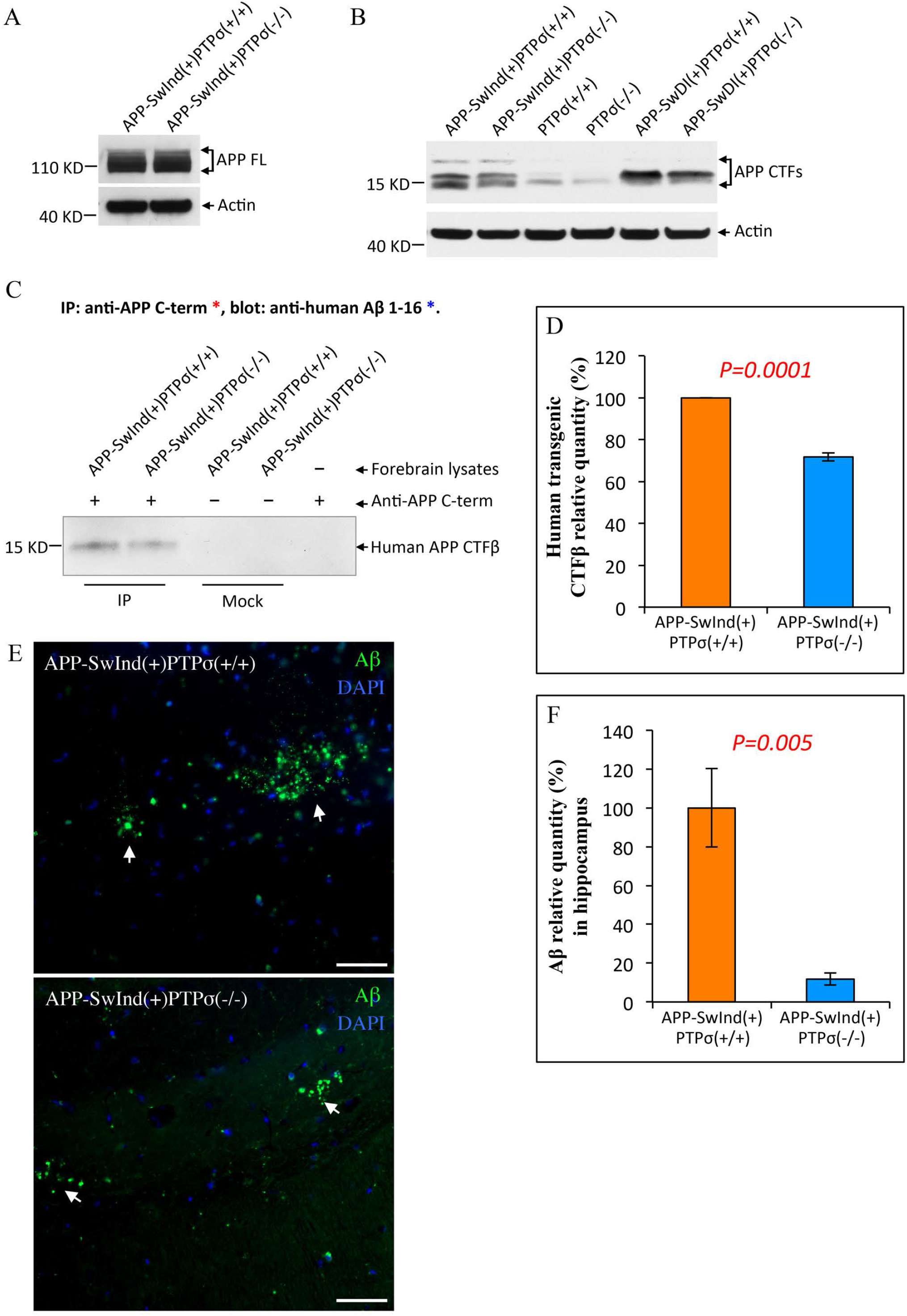
Genetic depletion of PTPσ reduces β-amyloidogenic products of APP. (**A**, **B**) Antibody (Y188) against the C-terminus of APP recognizes full length (FL) and C-terminal fragments (CTFs) of both mouse and human APP. PTPσ deficiency does not affect the expression level of APP FL (**A**), but reduces the level of an APP CTF at about 15 KD in mouse forebrain lysates (**B**). Images shown are representatives of at least three independent experiments. (**C**) Human CTFβ in the forebrains of APP-SwInd transgenic mice is identified using the method as described in Fig. 2D. CTFβ is immunoprecipitated by an antibody against the C-terminus of APP(Y188) and detected by western blot analysis using an antibody against amino acids 1-16 of human Aβ (6E10), which reacts with CTFβ but not CTFα (regions of antibody epitopes are shown in Fig. 2A). (**D**) Densitometry quantification of experiments as shown in panel **C** repeated with 5 pairs of gender-matched littermates. For each experiment, the value from PTPσ deficient sample was normalized to that from wild type sample. (**E**) Representative images of Aβ immunofluorescent staining (with 6E10) in the hippocampus of 15-month old TgAPP-SwInd mice. Arrows point to Aβ deposits. Scale bars, 50 µm. (**F**) ImageJ quantification of Aβ deposition in the hippocampus of 15-month old TgAPP-SwInd mice as shown in panel **E**. APP-SwInd(+)PTPσ(+/+), n=7; APP-SwInd(+)PTPσ(−/−), n=8. The mean value of APP-SwInd(+)PTPσ(−/−) samples was normalized to that of APP-SwInd(+)PTPσ(+/+) samples. All data are mean ± SEM. All *p* values, Student’s *t* test, 2-tailed.

**Figure S3.**
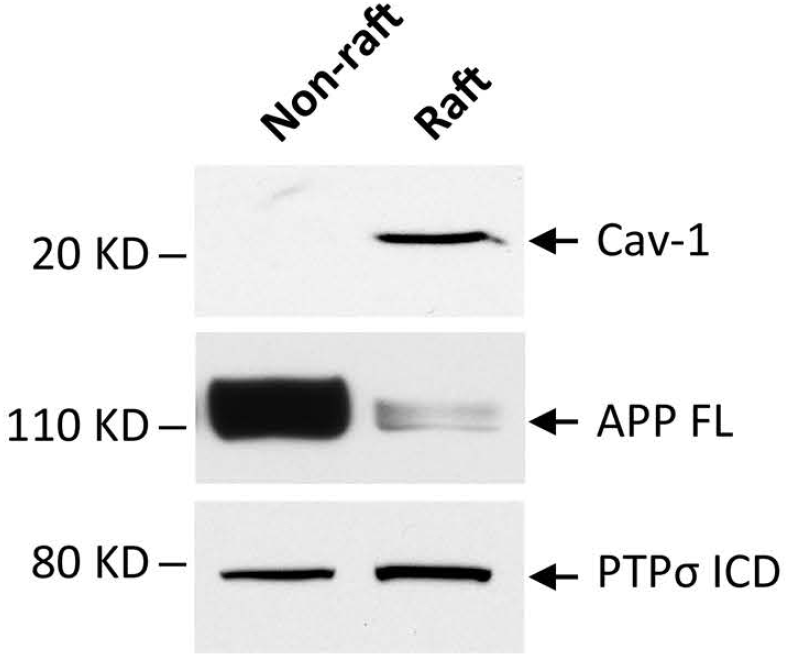
A small fractionation of APP is localized in lipid raft membrane domains. The lipid raft and non-raft membrane fractions were extracted from Balb/c mouse forebrains. Caveolin-1 (Cav-1), a scaffolding protein of caveola and a molecular marker of lipid rafts, is found only in the raft faction but not in the non-raft fraction. Full length APP (APP FL) is mostly distributed in the non-raft fraction with a significantly less amount in the lipid raft fraction, whereas PTPσ is found in both fractions. ICD, intracellular domain.

**Figure S4.**
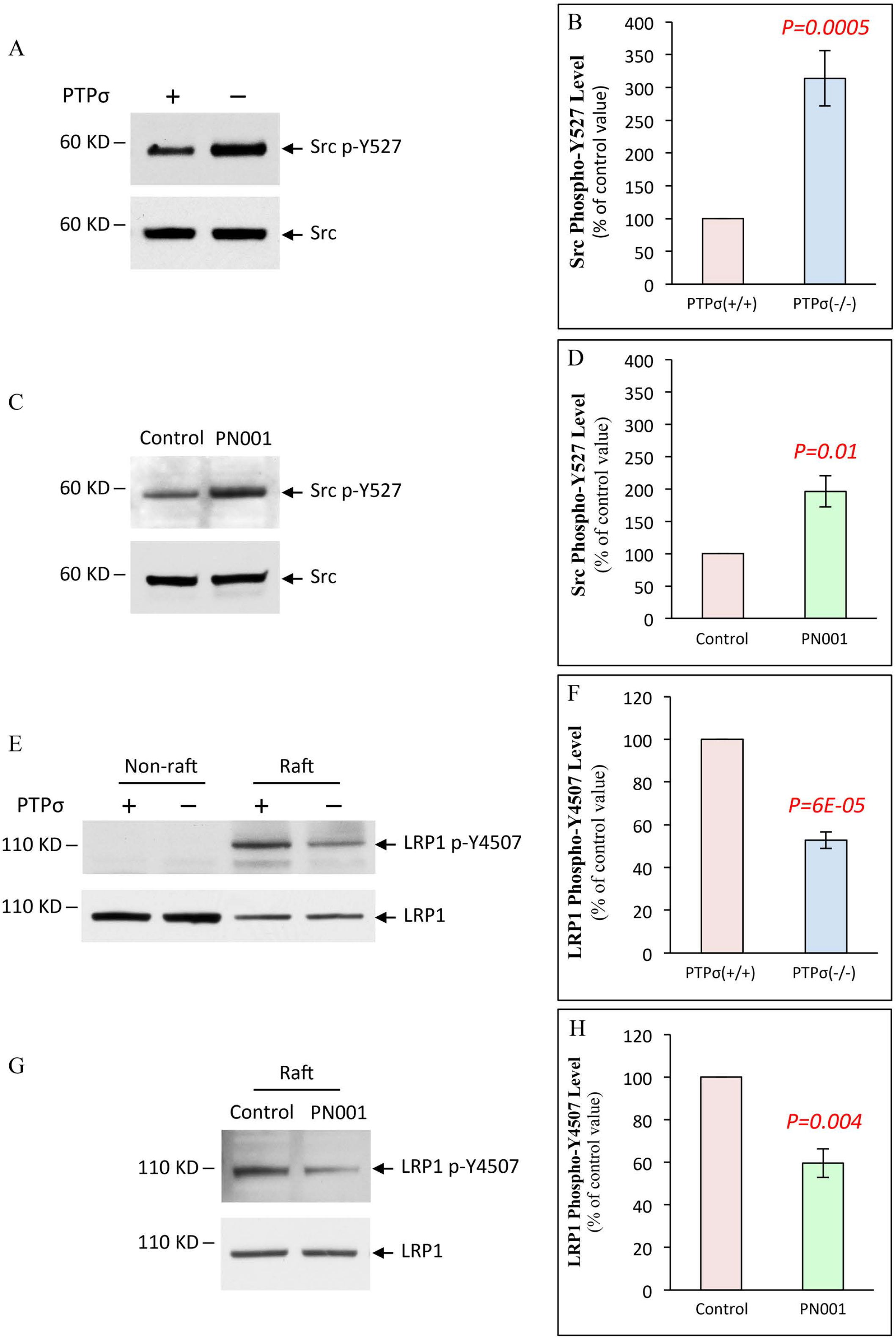
Modulators of APP *β*-processing are PTPσ downstream effectors. PTPσ genetic depletion (**A**, **B**, **E**, **F**) and pharmacological modulation by PN001 (**C**, **D**, **G**, **H**) result in similar effects on the tyrosine phosphorylation of Src kinase and its substrate LRP1, without affecting their expression levels. (**A**, **B**, **E**, **F**) Each experiment used a pair of gender-matched Balb/c littermates, with 1 wild type and 1 PTPσ-deficient. (**C**, **D**, **G**, **H**) Balb/c wild type littermates were treated via nasal administration of either vehicle solution (control) or PN001 (8 nmol), where the vehicle solution contains 5 nmol of γ-secretase inhibitor DAPT to prevent CTFβ degradation. Forebrains were collected 30 minutes later for analysis. Both methods of PTPσ downregulation lead to increased Src Y527 phosphorylation as shown by western blot analysis of forebrain lysates (**A**-**D**). LRP1 was analyzed in both non-raft and lipid raft membrane fractions. Y4507 phosphorylation of LRP1 localizes only in the lipid raft faction and its signal decreases in response to PTPσ downregulation (**E**-**H**). All LRP1 bands shown correspond to its C-terminal subunit. The non-raft and raft lanes were loaded with the same amount of total proteins. Results were quantified by densitometry. All data are mean ± SEM. *p* value, Student’s *t* test, 2-tailed. **A**, **B**, n=11; **C**, **D**, n=6; **E**, **F**, n=6; **G**, **H**, n=5.

**Figure S5.**
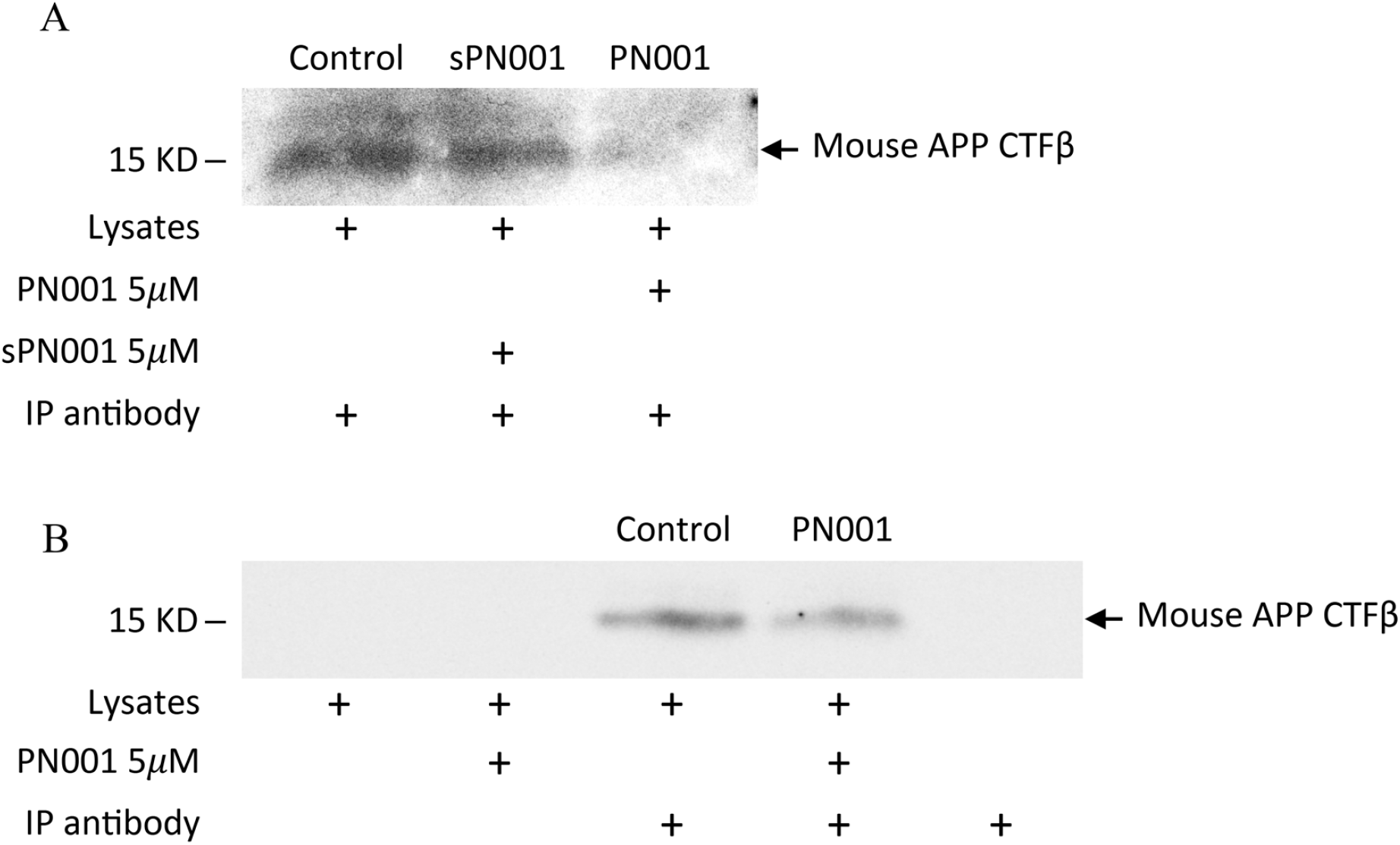
APP CTFβ level is dependent on PTPσ phosphatase activity. The effect of PTPσ inhibitory peptide PN001 on APP CTFβ level was assessed via *in vitro* (**A**) and *ex vivo* assays (**B**). (**A**) *In vitro* assays were performed using detergent free brain homogenates from Balb/c wild type mice that contain partially intact cellular membranes. Aliquots of crude brain homogenates were treated with the peptides as indicated at 37C° for 30 minutes with 1 µM of γ-secretase inhibitor DAPT to prevent CTFβ degradation. Control, vehicle, no peptide added. sPN001, a scrambled version of PN001. BACE1 is functional in the assay system as verified by the accumulation of APP CTFβ during the assay time frame (data not shown). (**B**) In *ex vivo* assays, the left and right hippocampal tissues were freshly isolated from Balb/c wild type mouse brain, and randomly chosen to be incubated with or without PN001 at 37C° for 30 minutes with 1 µM of DAPT. For both assays, at the end of incubation, membrane proteins were solubilized and mouse endogenous CTFβ was immunopurified using the same method as described in Fig. 2C. Mock immunoprecipitaion (IP) without antibody or lysates were used as negative controls. Each pilot assay was repeated (n=2) and representative images are shown. Treatment with PN001 reduces CTFβ level as compared to solution vehicle and sPN001. Further analyses and quantification of PN001 effects *in vivo* are shown in Fig. 5.

**Figure S6.**
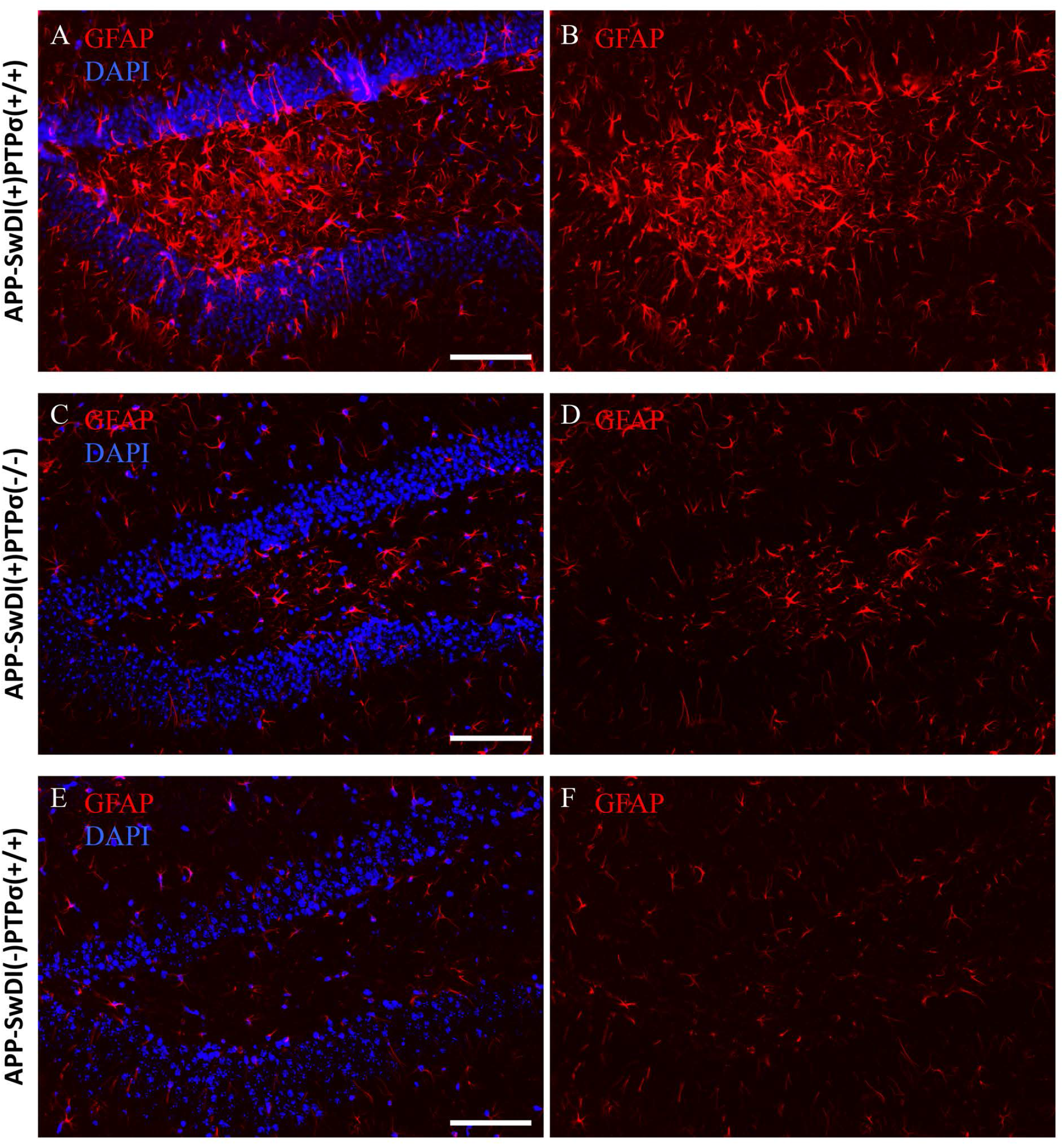
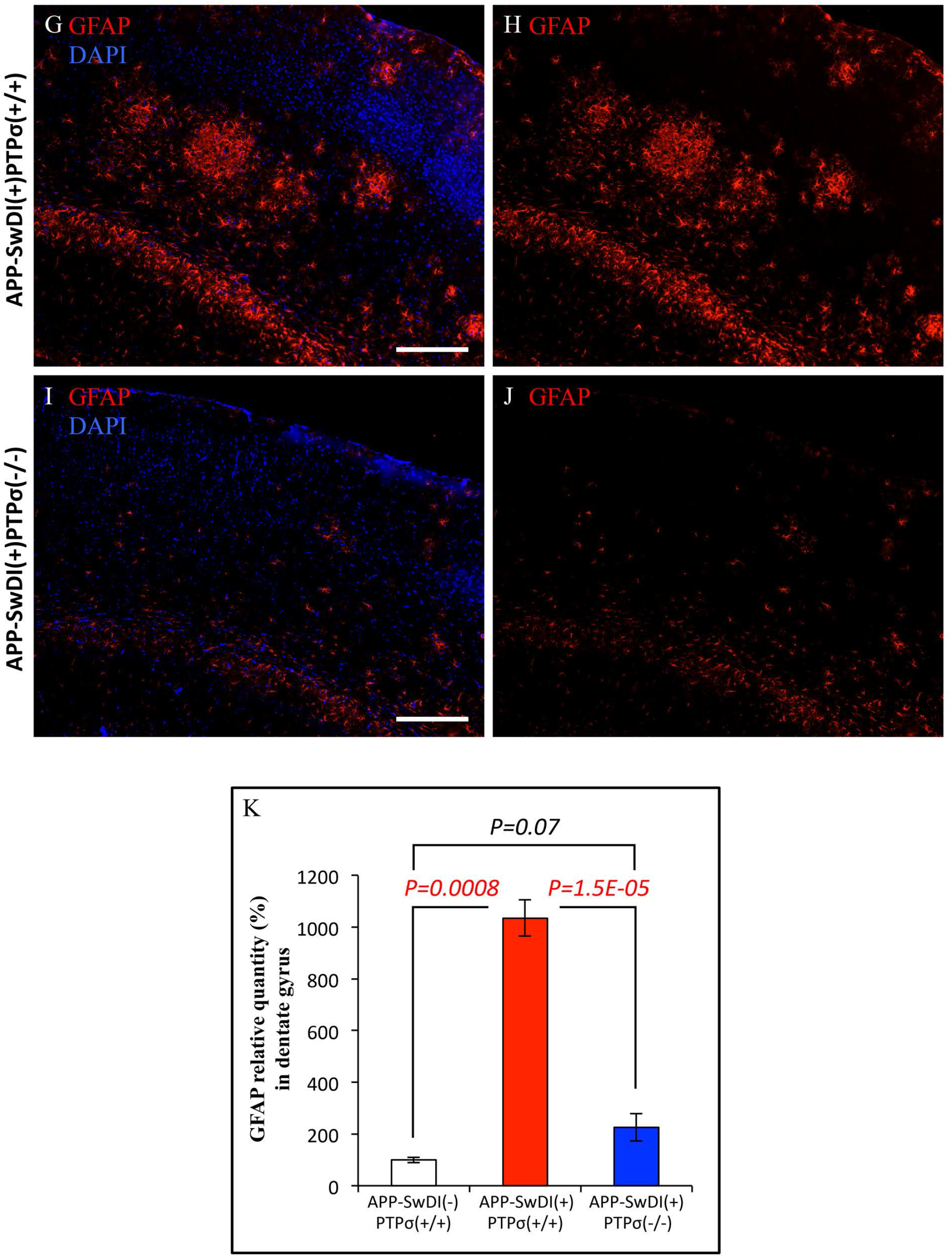
PTPσ deficiency attenuates reactive astrogliosis in APP transgenic mice. TgAPP-SwDI mice develop severe neuroinflammation as measured by the extortionate expression of GFAP, a marker of reactive astrocytes, which is restored close to normal level by PTPσ depletion in these mice. Representative images show GFAP (red) and DAPI staining of nuclei (blue) in the brains of 9-month old TgAPP-SwDI mice with or without PTPσ, along with their non-transgenic wild type littermates. (**A**-**F**) Dentate gyrus (DG) of the hippocampus; scale bars, 100 µm. (**G**-**J**) Primary somatosensory cortex; scale bars, 200 µm. (**K**) ImageJ quantification of GFAP level in DG hilus from mice aged between 9 to 11 months. APP-SwDI(-)PTPσ(+/+), non-transgenic wild type littermates. Total integrated density of GFAP in DG hilus was normalized to the area size of the hilus to yield average intensity as shown in the bar graph. Mean value of each group was normalized to that of APP-SwDI(-)PTPσ(+/+) mice. APP-SwDI(-)PTPσ(+/+), n=4; APP-SwDI(+)PTPσ(+/+), n=4; APP-SwDI(+)PTPσ(−/−), n=6. All *p* values, Student’s *t* test, 2-tailed. Data are mean ± SEM..

**Figure S7.**
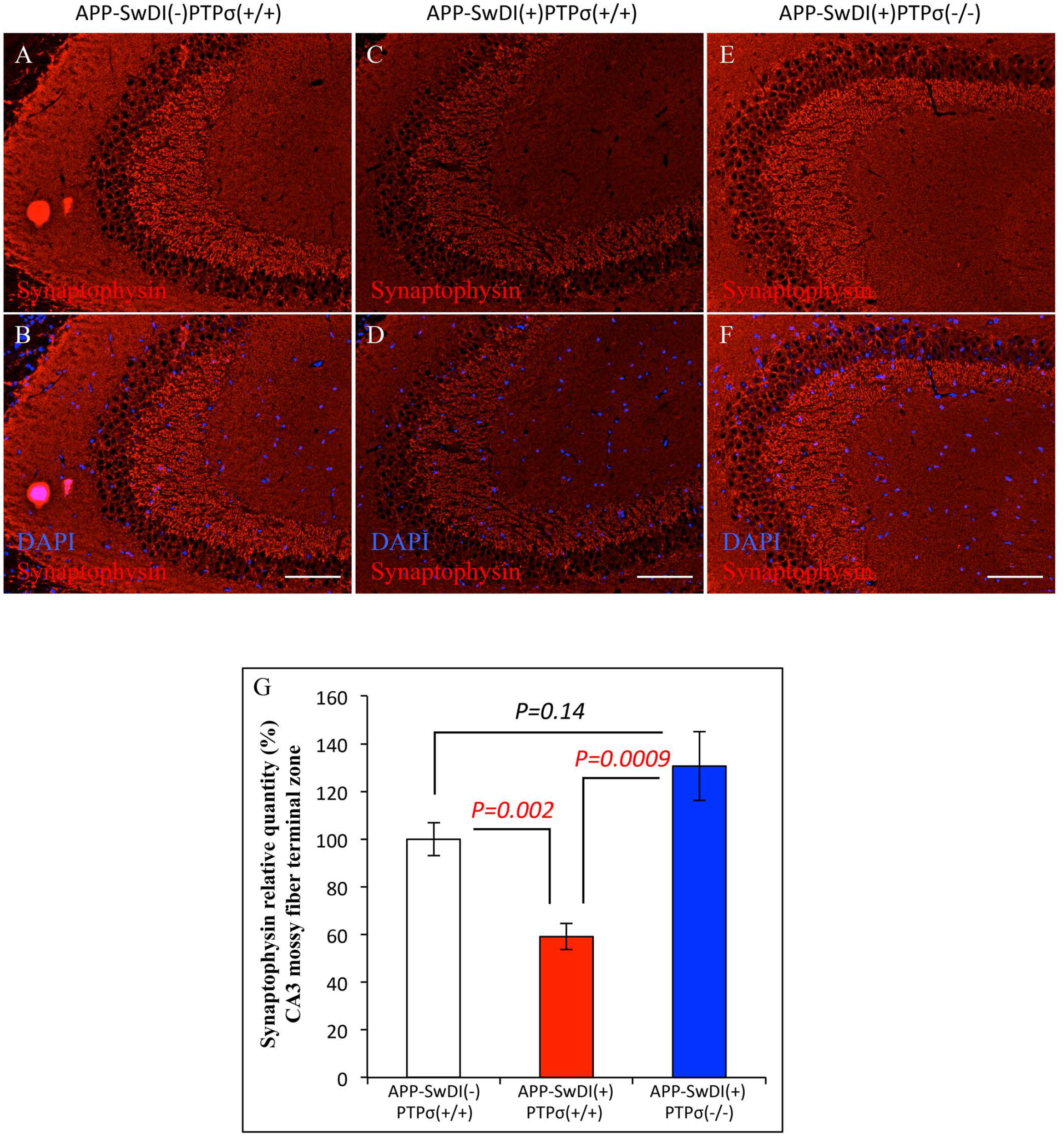
PTPσ deficiency protects APP transgenic mice from synaptic loss. Representative images show immunofluorescent staining of presynaptic marker Synaptophysin in the mossy fiber terminal zone of CA3 region. Synaptic loss is evident in TgAPP-SwDI mice, which is rescued by genetic depletion of PTPσ. (**A**-**F**) Synaptophysin, red; DAPI, blue. Scale bars, 100 µm. (**G**) ImageJ quantification of Synaptophysin expression level in CA3 mossy fiber terminal zone from mice aged between 9 to 11 months. Total integrated density of Synaptophysin in CA3 mossy fiber terminal zone was normalized to the area size to yield average intensity as shown in the bar graph. Mean value of each group was normalized to that of wild type APP-SwDI(-)PTPσ(+/+) mice. APP-SwDI(-)PTPσ(+/+), n=4; APP-SwDI(+)PTPσ(+/+), n=6; APP-SwDI(+)PTPσ(−/−), n=6. All *p* values, Student’s *t* test, 2-tailed. Data are mean ± SEM.

**Figure S8.**
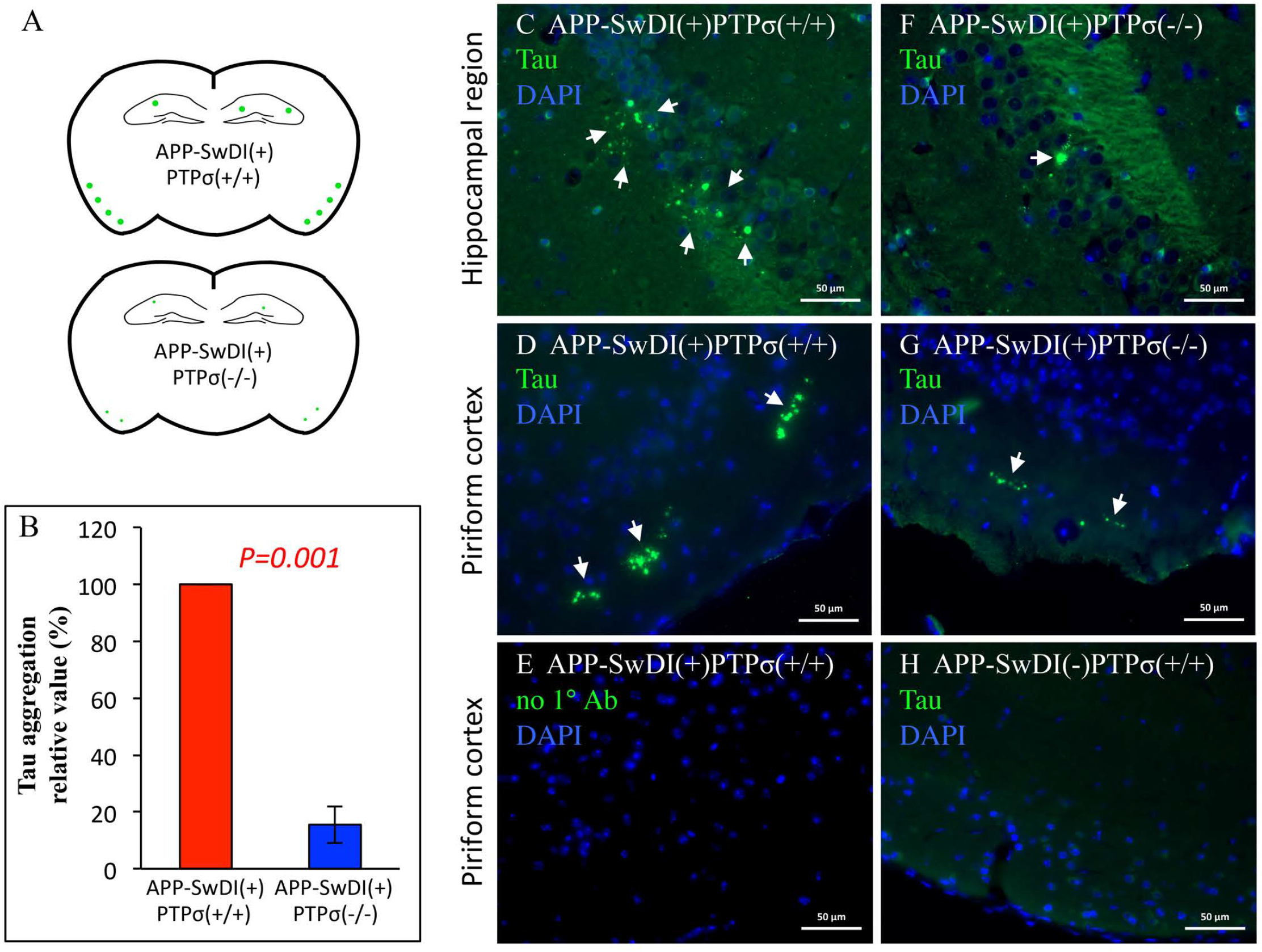
PTPσ deficiency mitigates Tau pathology in TgAPP-SwDI mice. (**A**) Schematic diagram depicting distribution pattern of Tau aggregation (green) detected by immunofluorescent staining, using an anti-Tau antibody (Tau-5) against its proline-rich region, in brains of 9 to 11 month-old TgAPP-SwDI transgenic mice. Similar results are seen with Tau-46, an antibody recognizing the C-terminus of Tau (Fig. S10I). Aggregated Tau is found most prominently in the molecular layer of piriform and entorhinal cortex, and occasionally in hippocampal regions in APP-SwDI(+)PTPσ(+/+) mice. (**B**) PTPσ deficiency diminishes Tau aggregation. Bar graph shows quantification of Tau aggregation in coronal brain sections from 4 pairs of gender-matched APP-SwDI(+)PTPσ(+/+) and APP-SwDI(+)PTPσ(−/−) littermates of 9 to 11 month-old. For each pair, the value from APP-SwDI(+)PTPσ(−/−) sample is normalized to the value from APP-SwDI(+)PTPσ(+/+) sample. *p* value, Student’s *t* test, 2-tailed. Data are mean ± SEM. (**C**, **D**) Representative images of many areas with Tau aggregation in APP-SwDI(+)PTPσ(+/+) brains. (**F**, **G**) Representative images of a few areas with Tau aggregation in brains of gender-matched APP-SwDI(+)PTPσ(−/−) littermates. **C** and **F**, Hippocampal regions. **D**-**H**, Piriform cortex. (**E**) Image shows the same area of Tau aggregation in **D** on an adjacent section stained without primary antibody (1° Ab). **H**, no Tau aggregates are detected in gender-matched non-transgenic wild type littermates. Tau, green; DAPI, blue. Arrows points to Tau aggregates. Scale bars, 50 µm.

**Figure S9.**
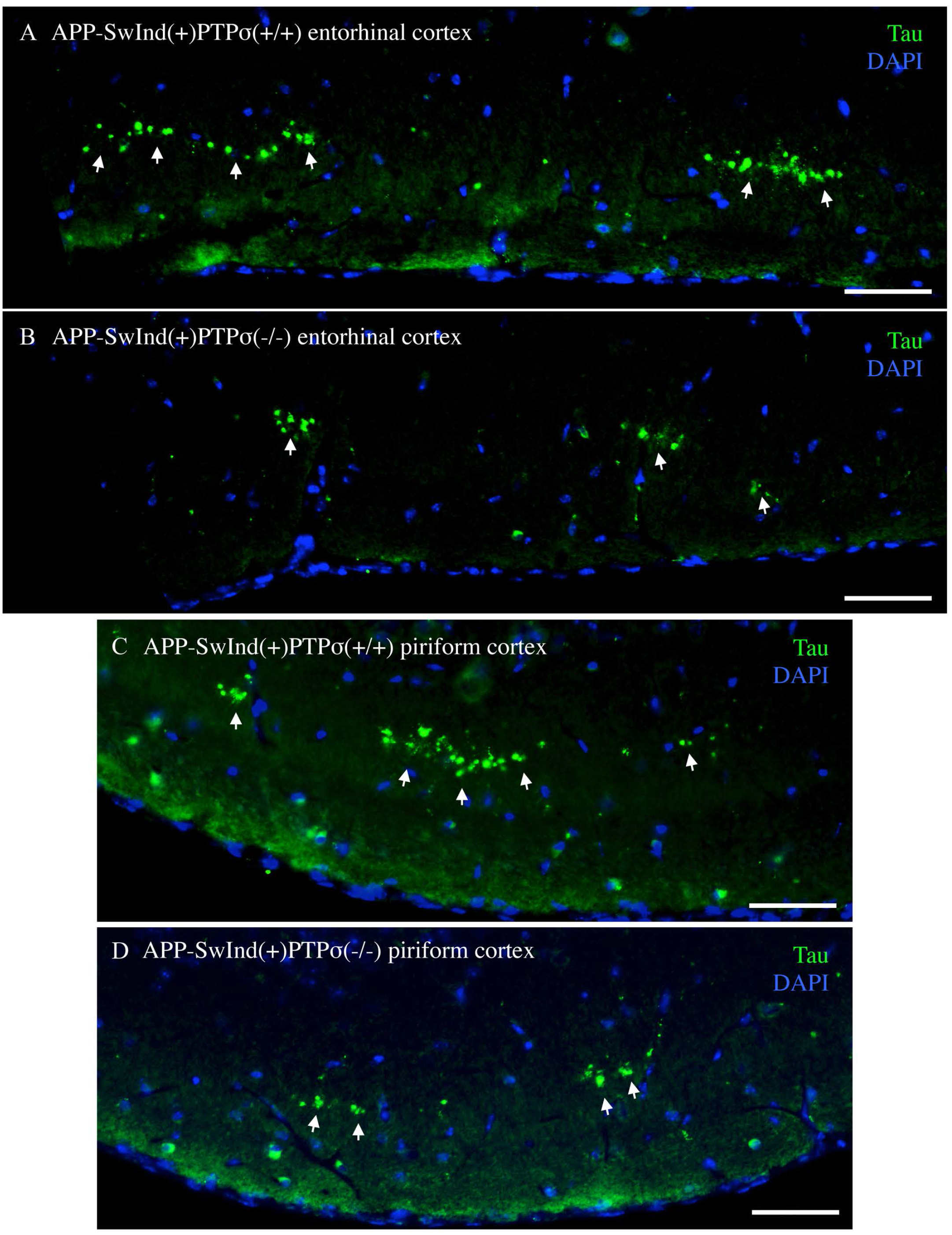
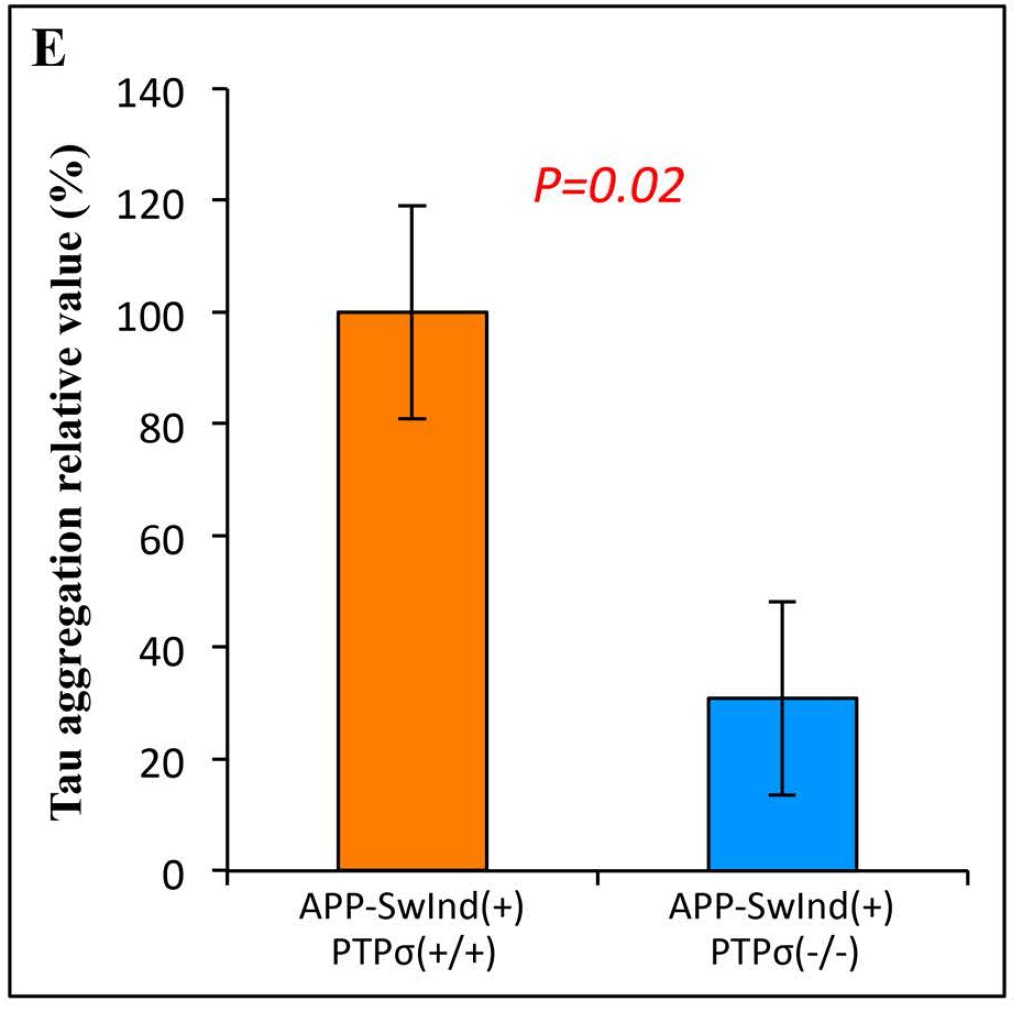
PTPσ deficiency mitigates Tau pathology in TgAPP-SwInd mice. Tau aggregation (green) is detected by immunofluorescent staining, using an anti-Tau antibody (Tau-5, as in Fig. 6 and Fig. S8) in the brains of 15 month-old TgAPP-SwInd transgenic mice. Similar results are observed with Tau-46, an antibody recognizing the C-terminus of Tau (data not shown). Aggregated Tau is found most prominently in the molecular layer of the entorhrinal (**A, B**) and piriform cortex (**C, D)**, and occasionally in the hippocampal regions (images not shown). (**E**) PTPσ deficiency diminishes Tau aggregation as quantified in coronal brain sections from 15 month-old APP-SwInd(+)PTPσ(+/+) (n=7) and APP-SwInd(+)PTPσ(−/−) mice (n=8). The mean value of APP-SwInd(+)PTPσ(−/−) samples is normalized to that of APP-SwInd(+)PTPσ(+/+). *p* value, Student’s *t* test, 2-tailed. Data are mean ± SEM. Tau, green; DAPI, blue. Arrows points to Tau aggregates. Scale bars, 50 µm.

**Figure S10.**
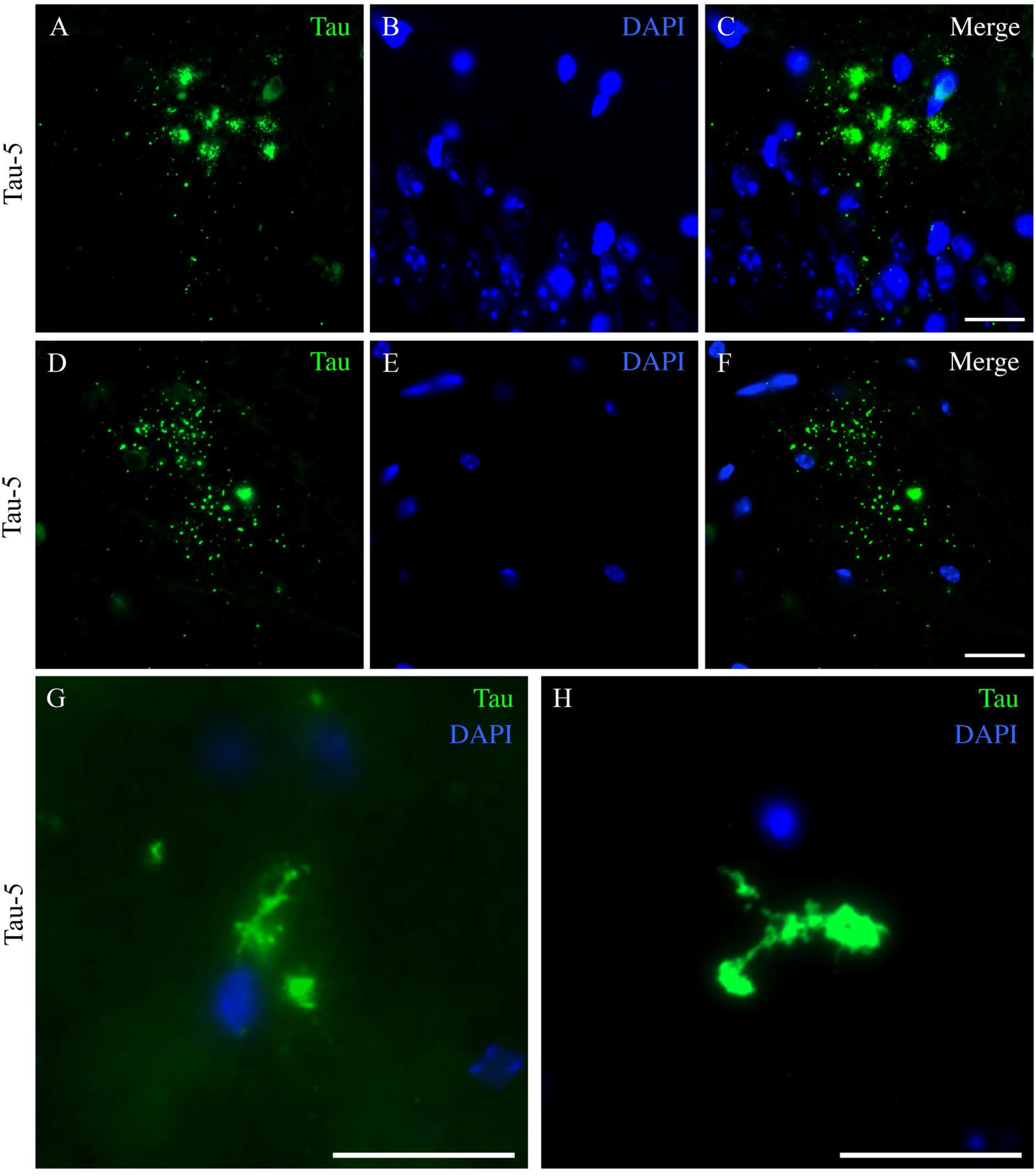
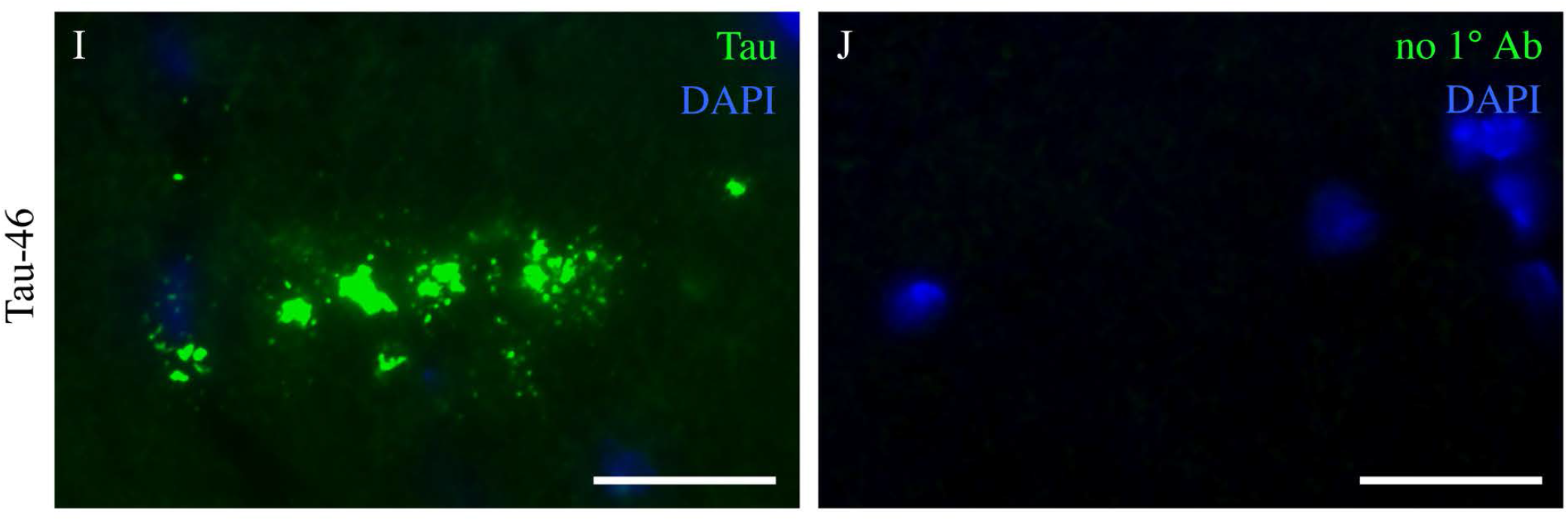
Morphology of Tau aggregates found in APP transgenic brains. (**A-H**) Tau aggregation (green) is detected by immunofluorescent staining, using an anti-Tau antibody (Tau-5) against the proline-rich domain of Tau (same as in Fig. 6, Fig. S8, and S9). Tau aggregates in TgAPP-SwDI and TgAPP-SwInd brains show similar morphologies. (**A-F**) Many of the Tau aggregates are found in punctate shapes, likely as part of cell debris, in areas that are free of nuclei staining. (**G, H**) Occasionally the aggregates are found in fibrillary structures, probably in degenerated cells before disassembling. (**I**) An additional anti-Tau antibody (Tau-46), which recognizes the C-terminus of Tau, detects Tau aggregation in the same pattern as Tau-5. (**J**) Image shows the same area of Tau aggregates in **I** on an adjacent section stained without primary antibody (1° Ab). Tau, green; DAPI, blue. All scale bars, 20 µm.

**Figure S11.**
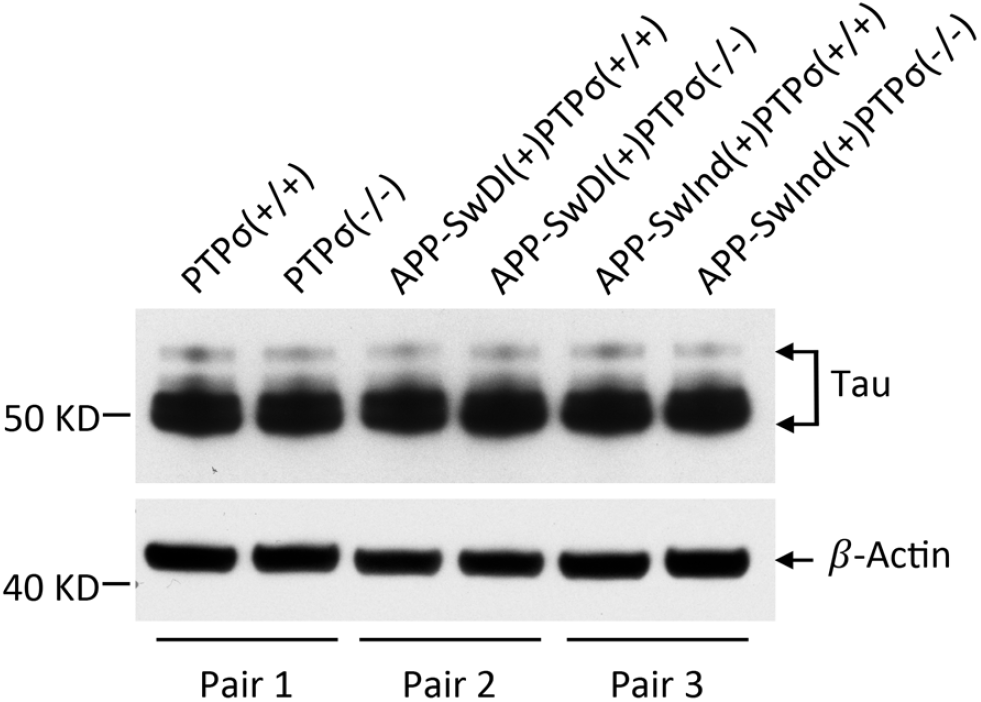
Tau expression is not affected by PTPσ or human APP transgenes. Upper panel, total Tau level in brain homogenates. Lower panel, *β*-Actin as loading control. Tau protein expression level is not changed by genetic depletion of PTPσ or expression of mutated human APP transgenes. All mice are older than 1 year, and mice in each pair are age- and gender-matched. Images shown are representatives of three independent experiments.

**Figure S12.**
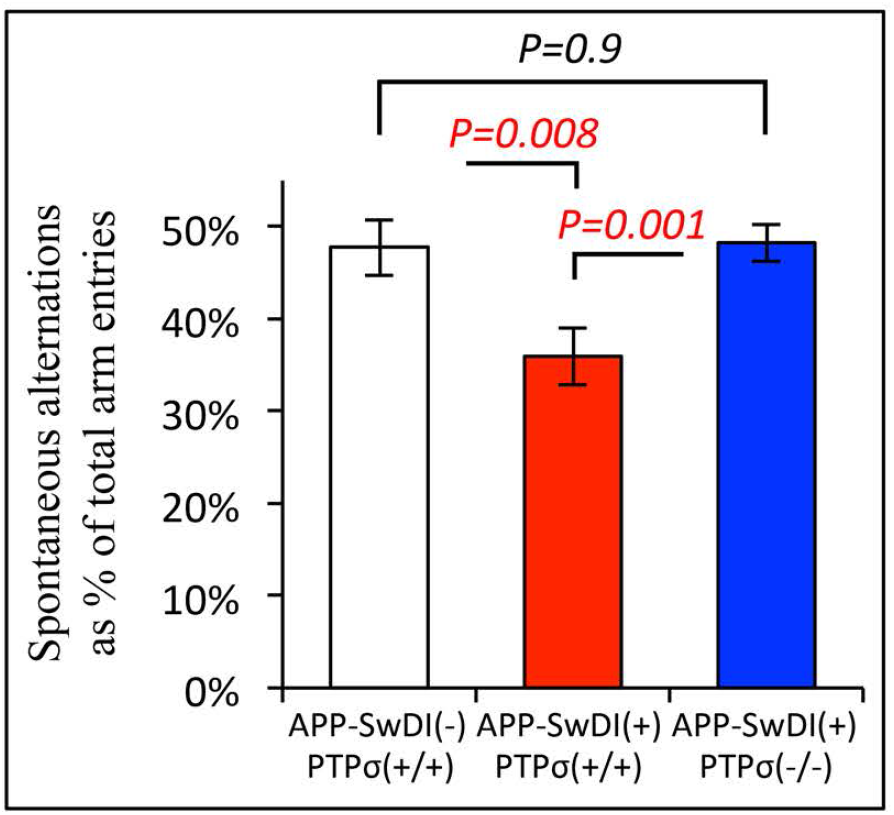
PTPσ deficiency restores short-term spatial memory in TgAPP-SwDI mice. In the Y-maze assay, performance of spatial navigation is scored by the percentage of spontaneous alternations among total arm entries. The raw values shown here are before normalization in Fig. 6M. Compared to non-transgenic wild type APP-SwDI(-)PTPσ(+/+)mice, APP-SwDI(+)PTPσ(+/+) mice show deficit of short-term spatial memory, which is rescued by genetic depletion of PTPσ. APP-SwDI(-)PTPσ(+/+), n=23 (18 females and 5 males); APP-SwDI(+)PTPσ(+/+), n=52 (30 females and 22 males); APP-SwDI(+)PTPσ(−/−), n=35 (22 females and 13 males). Ages of all genotype groups are similarly distributed between 4 and 11 months. All *p* values, Student’s *t* test, 2-tailed. Data are mean ± SEM.

**Figure S13.**
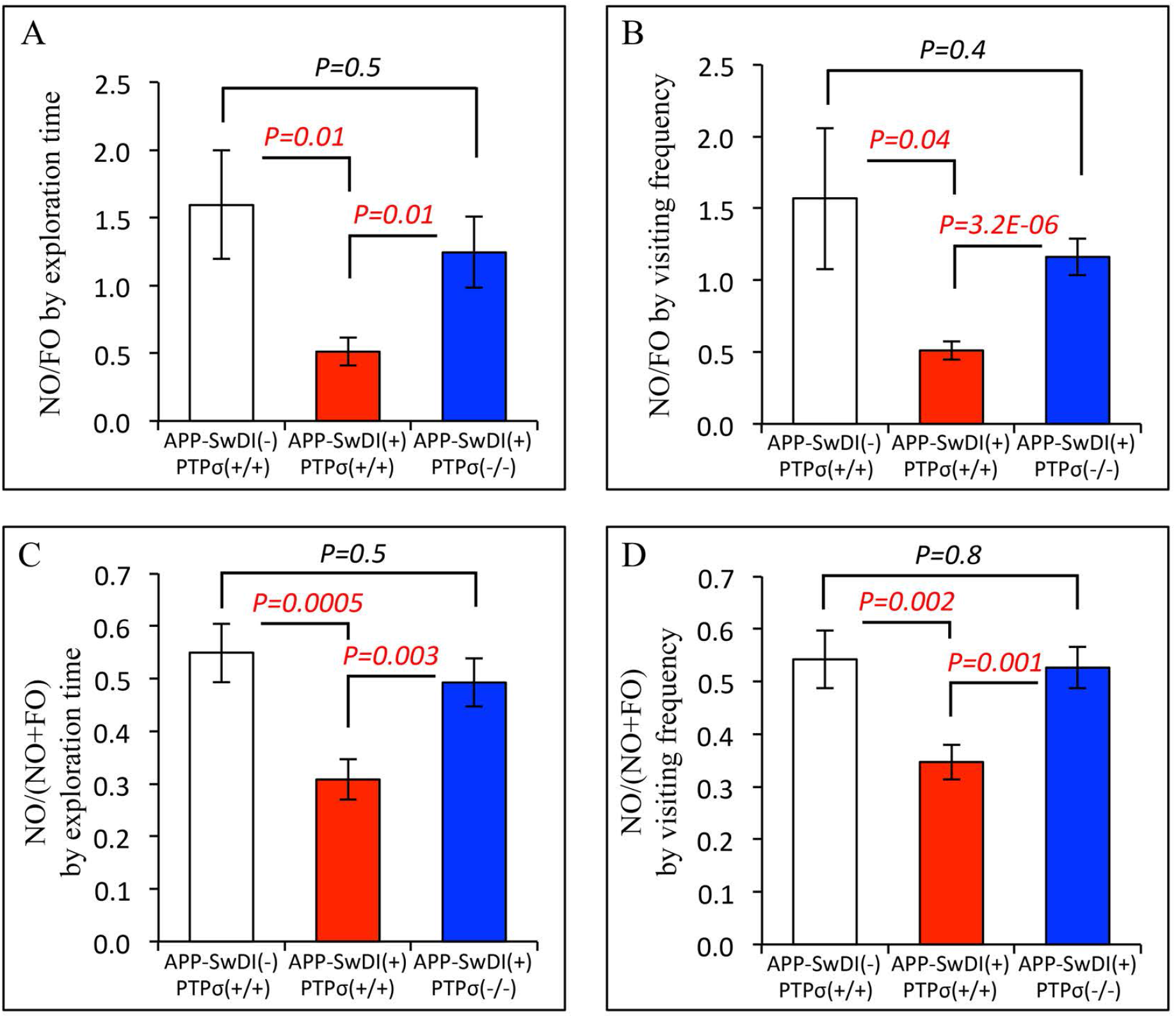
PTPσ deficiency enhances novelty exploration by TgAPP-SwDI mice. NO, novel object. FO, familiar object. (**A**, **B**) In novel object test, NO preference is measured by the ratio between NO and FO exploration, where NO/FO >1 indicates preference for NO. (**C**, **D**) Attention to NO is additionally measured by the discrimination index, NO/(NO+FO), the ratio of NO exploration to total object exploration (NO+FO). The raw values shown here in **C** and **D** are before normalization in Fig. 6N and O. Mice of this colony show a low baseline of the NO/(NO+FO) discrimination index, likely inherited from their parental Balb/c line. For non-transgenic wild type APP-SwDI(-)PTPσ(+/+) mice, the discrimination index is slightly above 0.5 (chance value), similar to the results previously reported for the Balb/c wild type mice (40). Thus, a sole measurement of the discrimination index may not reveal the preference for NO as does the NO/FO ratio. Although not as sensitive in measuring object preference, the NO/(NO+FO) index is most commonly used as it provides a normalization of the NO exploration to total object exploration activity. While each has its own advantage and shortcoming, both NO/FO and NO/NO+FO measurements consistently show that the expression of TgAPP-SwDI gene leads to a deficit in attention to the NO, whereas genetic depletion of PTPσ restores novelty exploration to a level close to that of non-transgenic wild type mice. **A** and **C**, measurements in terms of exploration time. **B** and **D**, measurements in terms of visiting frequency. APP-SwDI(-)PTPσ(+/+), n=28 (19 females and 9 males); APP-SwDI(+)PTPσ(+/ +), n=46 (32 females and 14 males); APP-SwDI(+)PTPσ(−/−), n=29 (21 females and 8 males). Ages of all groups are similarly distributed between 4 and 11 months. All *p* values, Student’s *t* test, 2-tailed. All data are mean ± SEM.

**Figure S14.**
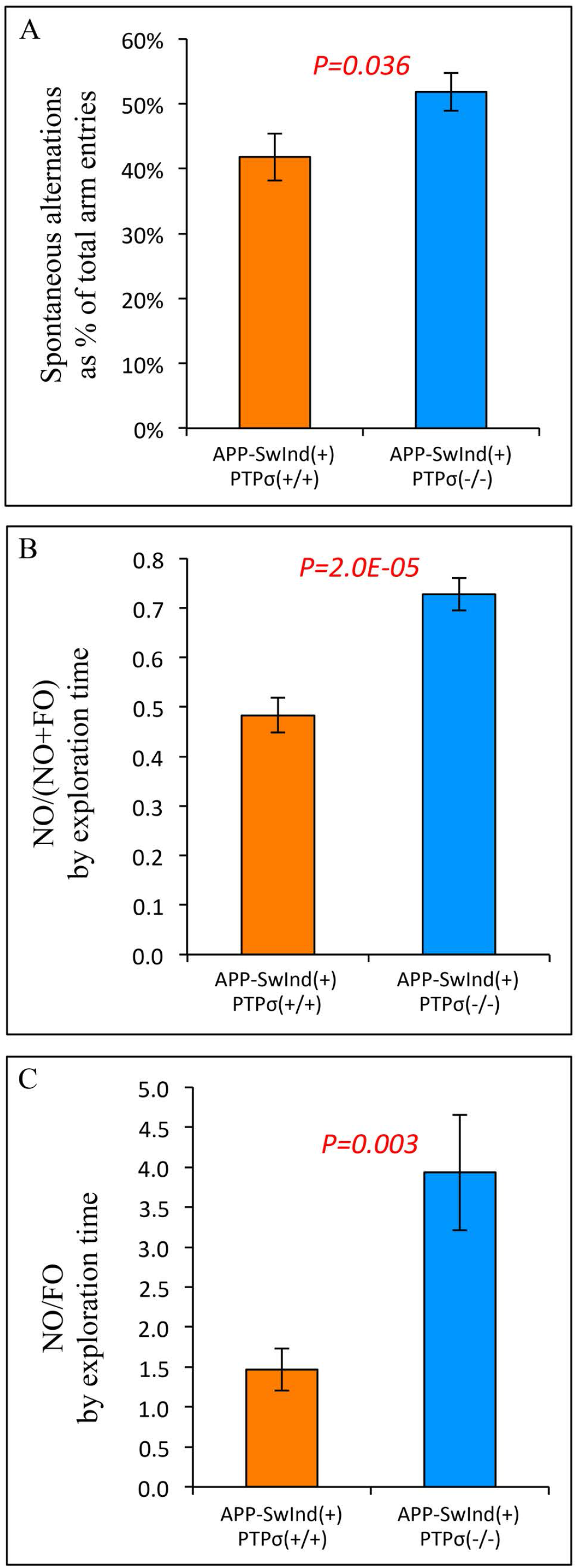
PTPσ deficiency improves behavioral performance of TgAPP-SwInd mice. (**A**) Performance of spatial navigation is scored by the percentage of spontaneous alternations among total arm entries in the Y-maze assay. Compared to APP-SwInd(+)PTPσ(+/+) mice, APP-SwInd(+)PTPσ(−/−) mice showed improved short-term spatial memory. APP-SwInd(+)PTPσ(+/+), n=40 (20 females and 20 males); APP-SwInd(+)PTPσ(−/−), n=18 (9 females and 9 males). Ages of both genotype groups are similarly distributed between 4 and 11 months. (**B, C**) Novel object test. NO, novel object. FO, familiar object. NO preference is measured by the ratio of NO exploration time to total object exploration time (**B**) and the ratio of NO exploration time to FO exploration time (**C**). PTPσ depletion significantly improves novelty preference in these transgenic mice. APP-SwInd(+)PTPσ(+/+), n=43 (21 females and 22 males); APP-SwInd(+)PTPσ(−/−), n=24 (10 females and 14 males). Ages of both groups are similarly distributed between 5 and 15 months. All *p* values, Student’s *t* test, 2-tailed. All data are mean ± SEM.

## Notes

### Competing Interest Statement

The authors have declared no competing interest.

### Summary of Updates

While the previous version showed data with PTPsigma genetic knockout, this new version has added pharmacological validation on the results.

